# Binary or non-binary fission? Reproductive mode of a predatory bacterium depends on prey size

**DOI:** 10.1101/2022.11.17.516866

**Authors:** Karolina Pląskowska, Łukasz Makowski, Agnieszka Strzałka, Jolanta Zakrzewska-Czerwińska

## Abstract

Most eukaryotic and bacterial cells divide by binary fission, where one mother cell produces two progeny cells, or, rarely, by non-binary fission. All bacteria studied to date use only one of these two reproduction modes. Here, we demonstrate for the first time that the predatory bacterium *Bdellovibrio bacteriovorus* reproduces through both binary and non-binary fission. Switching between the two modes correlates with the prey size. In relatively small host cells, *B. bacteriovorus* undergoes binary fission; the FtsZ ring assembles in the midcell and the mother cell splits into two daughter cells. In larger host cells, *B. bacteriovorus* switches to non-binary fission and creates multiple FtsZ rings to produce three or more daughter cells. Completion of bacterial cell cycle critically depends on precise spatiotemporal coordination of chromosome replication and segregation with other cell-cycle events, including cell division. Our studies reveal that *B. bacteriovorus* always initiates chromosome replication at the invasive pole of the cell, but the spatiotemporal choreography of subsequent steps depends on the fission mode and/or the number of progeny cells. In non-binary dividing filaments producing five or more progeny cells, the last round(s) of replication may also be initiated at the noninvasive pole. Finally, we show that binary-dividing *B. bacteriovorus* needs to extensively rebuild the flagellated pole of the mother cell to turn it into the invasive pole of a daughter cell. Altogether, we find that *B. bacteriovorus* reproduces through bimodal fission and that extracellular factors, such as the host size, can shape replication choreography, providing new insights about bacterial life cycles.

## Introduction

Most well studied model bacteria such as *Escherichia coli, Bacillus subtilis* and *Caulobacter crescentus,* reproduce by binary fission (1). In these bacteria, the newly synthesized sister chromosomes are segregated into two nascent cells prior to the completion of cell division. However, some bacteria belonging to various lineages, including antibiotic-producing *Streptomyces* and predatory *Bdellovibrio,* proliferate by non-binary fission (2). In such cases, more than two chromosome copies are synthesized and the resulting multinucleoid filamentous cell subdivides into single-nucleoid progeny cells (3). Thus, in growing non-binary bacteria, DNA replication is not directly followed by cell division.

While the dynamics of chromosome replication in relation to multi-point and synchronous septation have been relatively well studied in *Streptomyces* spp. (4–6), these aspects remain unexplored in *Bdellovibrio*. The most widespread species of this predatory genus is *Bdellovibrio bacteriovorus*, which preys on other Gram-negative bacteria. A growing body of research indicates that *B. bacteriovorus* could be used as a natural antibiotic agent (called a “living antibiotic”) in both healthcare and agriculture. *B. bacteriovorus* has been shown to kill various pathogens (e.g., *Klebsiella pneumoniae, Salmonella* and *Vibrio parahaemolyticus*) in different animal models (rats, mice, chicks and shrimps) (7–10).

In recent years, *B. bacteriovorus* has received considerable research attention, not only because of its potential application as a “living antibiotic”, but also because of its intriguing life cycle, which is reminiscent of that of a bacteriophage. The life cycle of *B. bacteriovorus* consists of two phases: a free-living non-replicative attack phase and an intracellular, non-binary reproductive phase (11–13). The free-living bacterium is asymmetric, with a flagellum and a pilus (i.e., an invasive pole) located at the opposite poles of cell. In the free-living phase, *B. bacteriovorus* uses its flagellum to move at speeds of up to 160 μm/s in search of prey (14). After engaging in pilus-mediated attachment to the prey’s outer membrane, *B. bacteriovorus* employs various glycosidases and peptidases to pierce the outer membrane and peptidoglycan, and then enters the periplasmic space (15, 16). At the beginning of the reproductive phase, the host cell dies and bloats into a spherical structure called a prey bdelloplast, inside of which the predatory cell “consumes” the prey contents (i.e., it degrades the prey’s macromolecules and reuses them for its own growth) and proliferates by non-binary fission. During this phase, the single chromosome is copied multiple times. Chromosome replication is initiated near the invasive pole (17, 18) and, after a few rounds of chromosome replication, the resulting multinucleoid filamentous cell divides into various (even or odd) numbers of progeny cells. Usually three to six *B. bacteriovorus* progeny cells are formed when *E. coli* serves as the host (19). The flagellated progeny cells are released into the environment to repeat the cycle.

Our current knowledge of the *B. bacteriovorus* cell cycle is largely based on studies employing *E. coli* as a model host. In order to understand how *B. bacteriovorus* can be applied to combat bacterial infections caused by pathogens, studies on its life cycle in different Gram-negative pathogenic bacteria are of paramount importance.

In the present study, we provide key insights into the mode and dynamics by which *B. bacteriovorus* proliferates in different pathogens, *Proteus mirabilis, Salmonella enteritidis* and *Shigella flexneri* that are on the World Health Organization (WHO) global priority pathogen list of antibiotic-resistant bacteria (20). Currently, at least 700,000 people worldwide die each year due to drug-resistance infections and according to the WHO predictions, this number could rise to 10 million by 2050 (21). Using a set of *B. bacteriovorus* fluorescent reporter strains, we performed real-time observation of the major steps of the predator’s cell cycle in different pathogens. These steps include chromosome replication and cell division (septation). We reveal that the chromosome replication choreography and division mode differs across the analyzed host pathogens, and demonstrate for the first time that the predatory bacterium, *B. bacteriovorus,* undergoes binary or non-binary fission depending on the size of its host cell.

## Results

### Proliferation of *B. bacteriovorus* in different pathogens

To investigate the influence of the prey cell on *B. bacteriovorus* proliferation, we chose three different hosts that are on the WHO global priority pathogen list of antibiotic-resistant bacteria (20): *Proteus mirabilis, Salmonella enteritidis* and *Shigella flexneri* (for details see Table S1). Firstly, we assessed the ability of *B. bacteriovorus* to prey on the different pathogens, by generating predatory killing curves using the pathogens as hosts (Fig. S1A). In this analysis, a decrease in prey cell optical density reflected the lysis of cells by *B. bacteriovorus*. Interestingly, the shapes of the curves are different among the analyzed hosts. The fastest decrease in prey cell optical density was observed for *S. flexneri* cells. Similar dynamics of predation was observed for *S. enteritidis*. While the dynamics of *B. bacteriovorus* predation on the third tested pathogen *P. mirabilis* cells was substantially different: there was slow decrease in optical density at the beginning of measurements followed by a rapid decrease in optical density (Fig. S1A).

To further elucidate the differences in predation dynamics (mirrored by the killing curves), we measured different parameters of the prey and predator cells, including the host cell size before infection, the diameter of the formed bdelloplasts (Fig. S1B) and the number of predatory cells released after host cell lysis (Fig. 1) using single cell microscopic data. The average size of the host cell varied and was reflected by differences in the bdelloplast diameter (Fig. S1B). As expected, the diameter of the bdelloplast was positively correlated with the length of the prey cell (calculated Pearson correlation coefficient R=0.87). The number of predatory cells formed inside the bdelloplast positively correlated with the length of the host cell (Fig. 1). Surprisingly, predation on a *P. mirabilis* cell resulted consistently in the generation of two progeny cells (100%, n=100 prey cells analyzed; Fig. 1). On the contrary more progeny cells were formed in larger-sized hosts: two to five (usually 3-4) and four to eight (usually 5-7) progeny cells were produced from *S. enteritidis* and *S. flexneri,* respectively (Fig. 1). Thus, the infection dynamics of *B. bacteriovorus* in a host cell species is related to the number of progeny cells released from a single cell of that host species. The predatory killing curves illustrated this correlation (Fig. S1A): At the beginning of infection, when more progeny predators were released (e.g., from *S. flexneri* cells) more prey cells were killed, as reflected by a faster decrease in the prey cell optical density (Fig. S1A).

**Figure 1.**
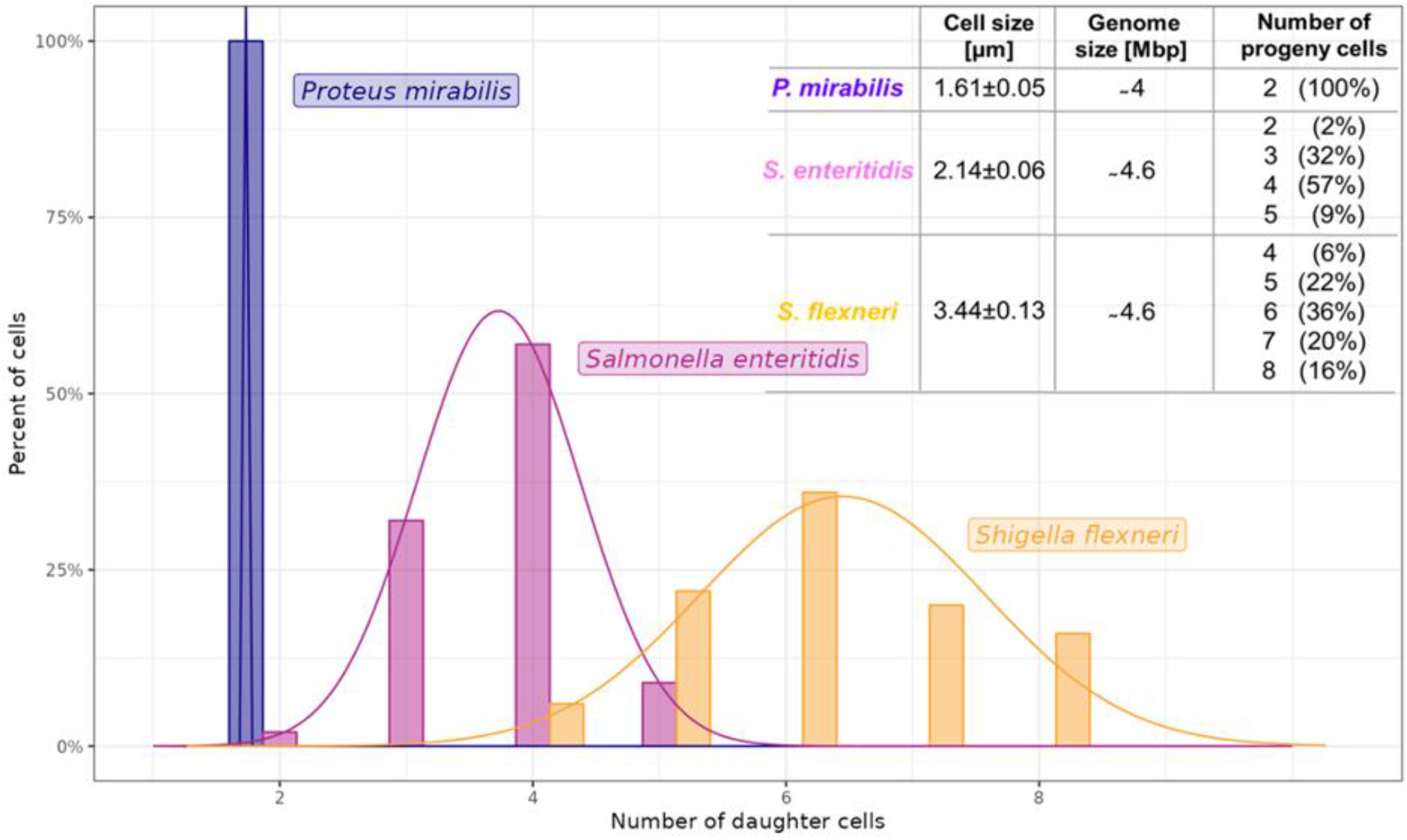
Number of progeny cells of *B. bacteriovorus* formed in different hosts (n=100 for each host).

In sum, our results suggest for the first time that *B. bacteriovorus* divides not only by non-binary (more than two even or odd daughter cells), but also by binary (two daughter cells in *P. mirabilis)* fission, and the mode of division appears to depend on the host size.

### Life cycle of *B. bacteriovorus* in different pathogens

As we observed host-dependent differences in predation dynamics, we next decided to closely investigate the main stages of the *B. bacteriovorus* reproductive phase (i.e., chromosome replication, segregation and filament septation) in *P. mirabilis*, *S. enteritidis* and *S. flexneri* at the single-cell level using time-lapse fluorescence microscopy (TLFM). For this purpose, we constructed a set of *B. bacteriovorus* reporter strains that allowed us to monitor the localization of the replisome (*B. bacteriovorus* DnaN-mNeonGreen) or simultaneously assess the localizations of the replisome and segrosome (*B. bacteriovorus* mNeonGreen-ParB/DnaN-mCherry) or the replisome and divisome *B. bacteriovorus* (*B. bacteriovorus* FtsZ-mNeonGreen/DnaN-mCherry). The characteristic features of the analyzed proteins (DnaN, ParB and FtsZ) are presented in Table 1. In all constructed strains, the gene of interest (*dnaN, parB* or *ftsZ*) was exchanged for a fusion gene that was located in the native chromosomal locus and expressed under its own native promoter(s) (see Table S1). Production of the fusion proteins was confirmed using microscopic analysis (Figs. 2–7, S5 and S7-S9) or Western blotting (Fig. S3). *B. bacteriovorus* reporter strains exhibited a killing curve for *E. coli* (Fig. S4) similar to that of the wild-type strain.

**Table 1.**
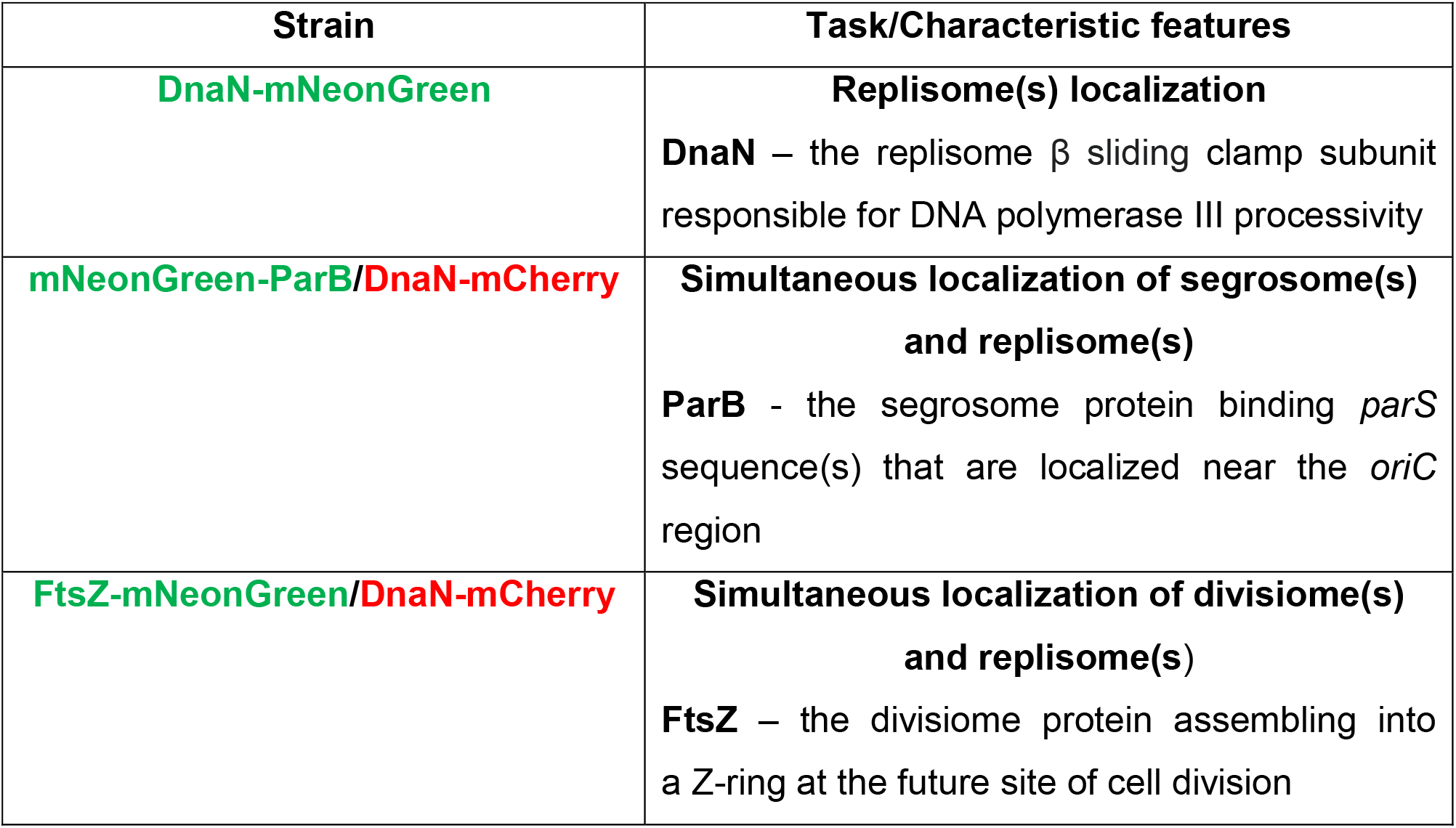
Characteristic features of the *B. bacteriovorus* fluorescent reporter strains used in this study.

| Strain | Task/Characteristic features |
| --- | --- |
| **DnaN-mNeonGreen** | **Replisome(s) localization**<br>**DnaN** – the replisome $\beta$ sliding clamp subunit responsible for DNA polymerase III processivity |
| **mNeonGreen-ParB/DnaN-mCherry** | **Simultaneous localization of segrosome(s) and replisome(s)**<br>**ParB** - the segrosome protein binding *parS* sequence(s) that are localized near the *oriC* region |
| **FtsZ-mNeonGreen/DnaN-mCherry** | **Simultaneous localization of divisiome(s) and replisome(s)**<br>**FtsZ** – the divisiome protein assembling into a Z-ring at the future site of cell division |

**Figure 2.**
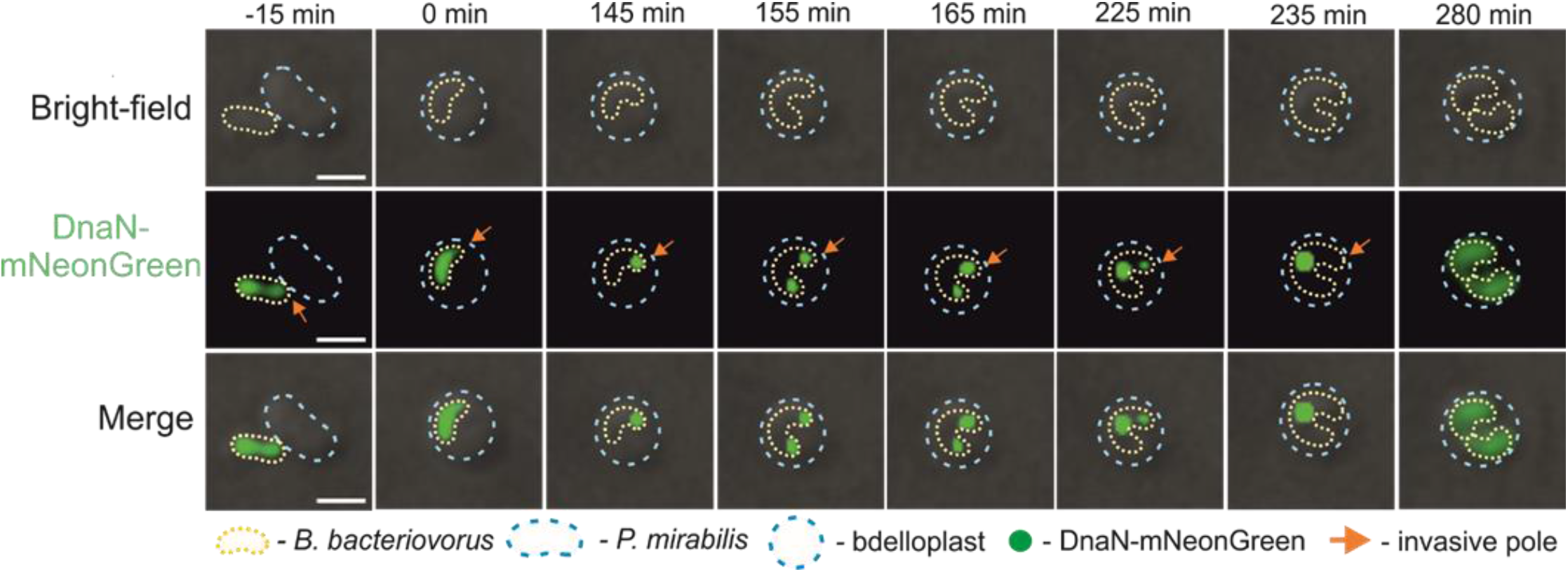
Spatiotemporal analysis of chromosome replication in a *B. bacteriovorus* cell growing in *P. mirabilis*. Time-lapse analysis of a representative *B. bacteriovorus* cell showing the localization of replisome(s) (green) in a predatory cell growing inside the *P. mirabilis* bdelloplast. (**A**) Attachment of *B. bacteriovorus* to a *P. mirabilis* cell. (**B**) Bdelloplast formation, time = 0 min. (**C**) Appearance of the first DnaN-mNeonGreen focus at the invasive pole of a *B. bacteriovorus* cell, indicating initiation of chromosome replication. (**D**-**G**) Further steps of chromosome replication, including splitting of replication forks (the appearance of the second DnaN-mNeonGreen focus at the “flagellar” pole) and merging of replication forks at the midcell. (**H**) Termination of chromosome replication (disassembly of replisomes) and formation of two progeny cells. Photos represent merged bright-field and green fluorescence images. Scale bar = 1 μm. The full time-lapse is shown in Video S1.

**Figure 3.**
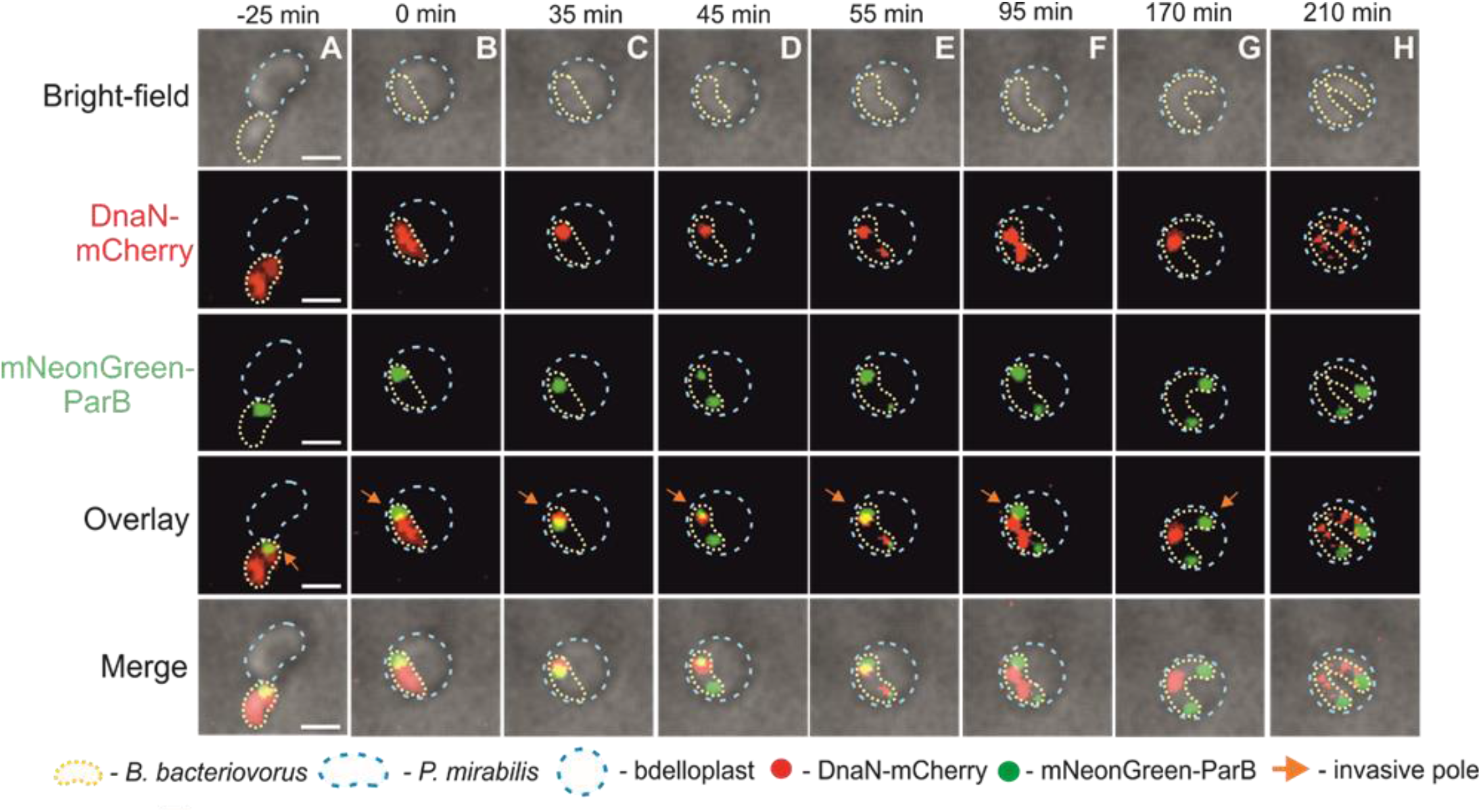
Time-lapse analysis of chromosome replication and *oriC* segregation in a *B. bacteriovorus* cell growing in *P. mirabilis*. Localization of replisome(s) (red) and *oriC* (i.e., ParB complex, green) in a predatory *B. bacteriovorus* cell growing inside a *P. mirabilis* bdelloplast. (**A**) Attachment of *B. bacteriovorus* to a *P. mirabilis* cell. (**B**) Bdelloplast formation, time = 0 min. (**C**) Appearance of the first DnaN-mCherry focus at the invasive pole of a *B. bacteriovorus* cell, indicating initiation of chromosome replication. (**D-G**) Duplication of the *oriC* region (i.e., mNeonGreen-ParB focus) at the invasive pole; thereafter, one of the two *oriC* regions migrates to the opposite pole and remains there until the end of the cell cycle. Further steps of chromosome replication include splitting of replication forks (the appearance of the second DnaN-mCherry focus at the flagellar pole) and merging of replication forks at the midcell. (**H**) Termination of chromosome replication (disassembly of replisomes) and formation of two progeny cells. Photos represent merged bright-field and fluorescence (red and green) images. Scale bar = 1 μm. The full time-lapse is shown in Video S2.

**Figure 4.**
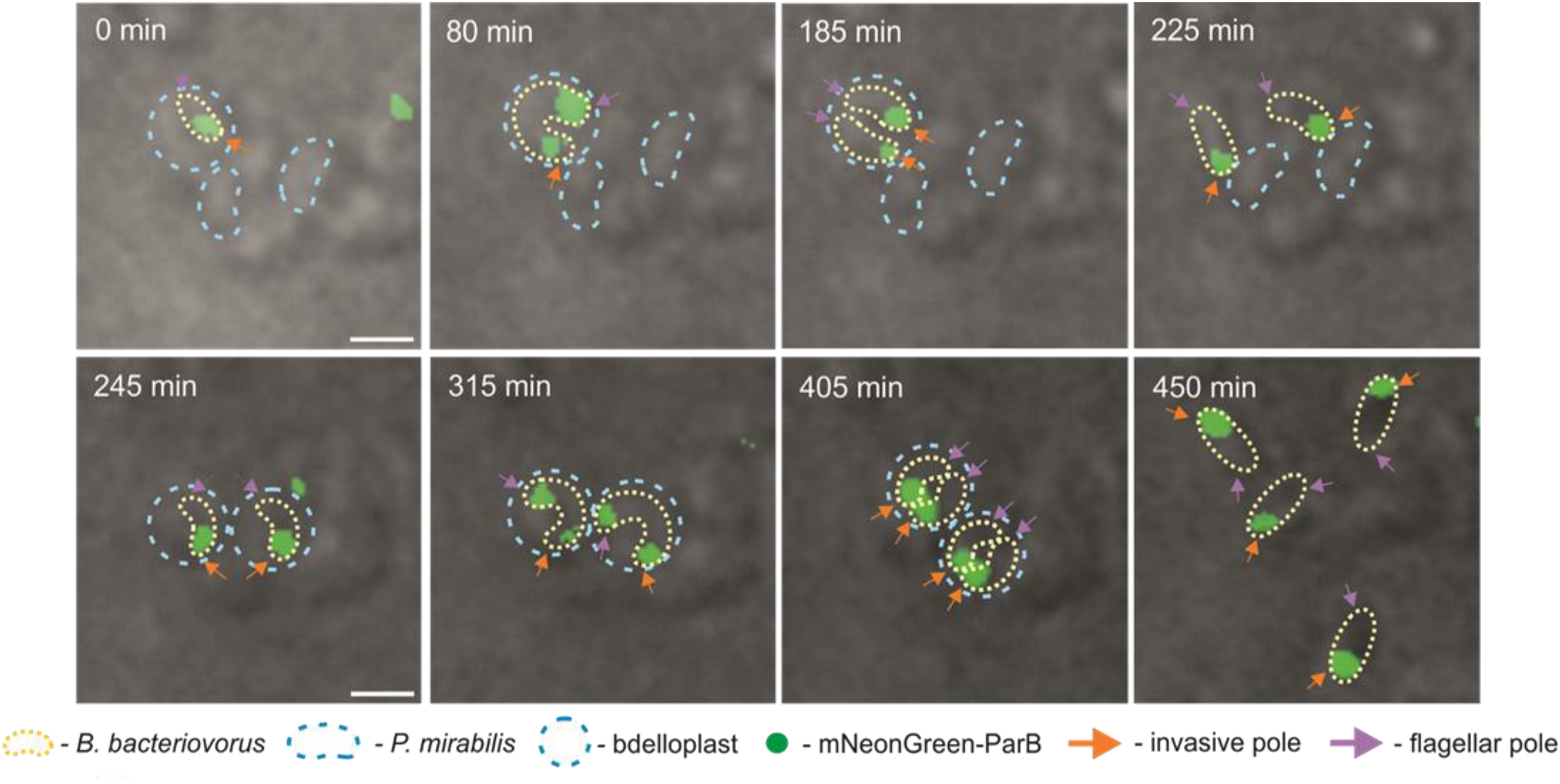
Conversion of the flagellated pole of the mother cell into an invasive pole inherited by a daughter cell during *B. bacteriovorus* proliferation in *P. mirabilis*. Time-lapse microscopy of *B. bacteriovorus* growing inside *P. mirabilis* shows that the flagellated pole of the mother cell (violet arrow) becomes an invasive pole (orange arrow) that is inherited by the daughter cell. The newly formed invasive pole, has a visible ParB complex (i.e., *oriC* region) and is functional, since the daughter cell attacks the next *P. mirabilis* cell through that pole. Photos represent merged bright-field images. The *B. bacteriovorus* cell and the bdelloplast are marked by yellow and blue dotted lines, respectively. Scale bar = 1 μm. The full time-lapse is shown in Video S3.

**Figure 5.**
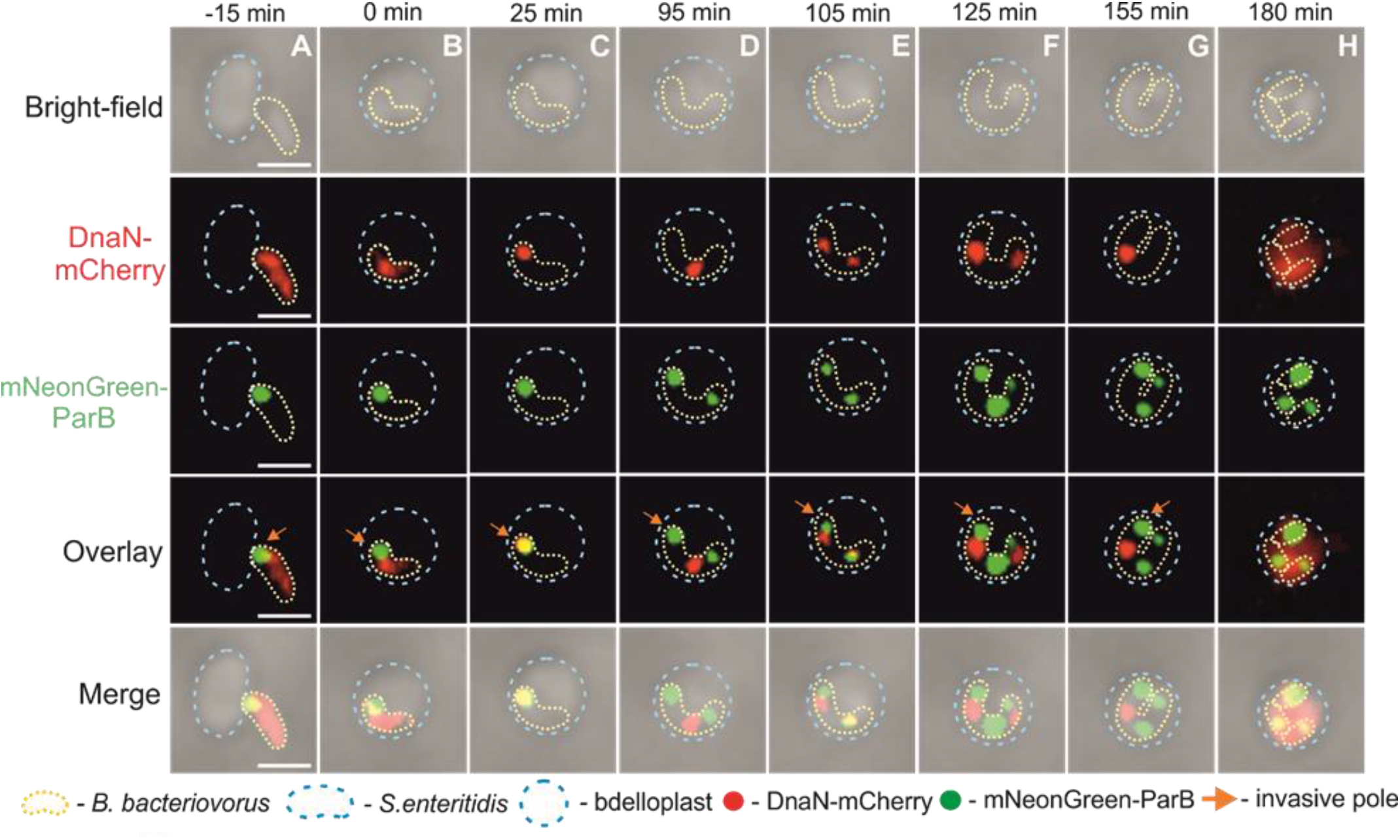
Dynamics of chromosome replication and *oriC* segregation in a *B. bacteriovorus* cell growing in *S. enteritidis: formation of three daughter cells*. (**A**) Attachment of *B. bacteriovorus* to an *S. enteritidis* cell. (**B**) Bdelloplast formation, time = 0 min. (**C**) Appearance of the first DnaN-mCherry focus at the invasive pole of a *B. bacteriovorus* cell, indicating initiation of chromosome replication. (**D**) Duplication of the *oriC* region (i.e., mNeonGreen-ParB focus) at the invasive pole; thereafter, one of the two *oriC* regions migrates to the opposite pole and one of the two DnaN-mCherry foci follows behind the newly replicated *oriC* region that is already attached to the opposite pole. (**E and F**) Reinitiation of DNA replication from the mother chromosome located at the invasive pole, migration of both replisomes toward the opposite pole and the appearance of the third ParB focus at the middle of the filament. (**G**) Disappearance of the DnaN-mCherry focus located near the flagellar pole, indicating termination of the first round DNA replication. (**H**) Termination of the second round of chromosome replication and formation of three daughter cells. Photos represent merged bright-field and fluorescence (red and green) images. The *B. bacteriovorus* cell and the bdelloplast are marked by yellow and blue dotted lines, respectively. Scale bar = 1 μm. The full time-lapse is shown in Video S4.

**Figure 6.**
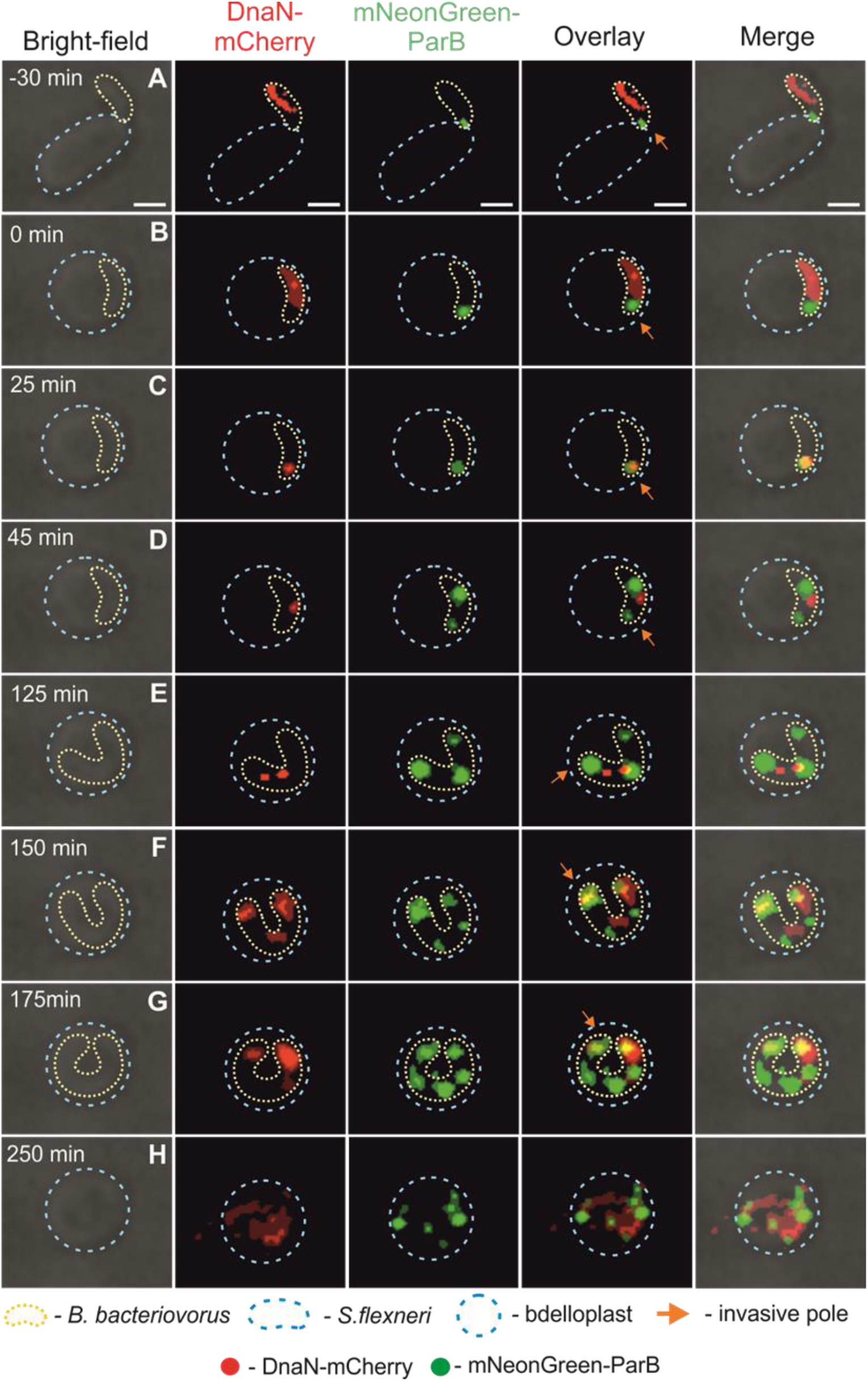
Dynamics of chromosome replication and *oriC* segregation in a *B. bacteriovorus* cell during proliferation in *S. flexneri*. (**A**) Attachment of *B. bacteriovorus* to an *S. flexneri* cell. (**B**) Bdelloplast formation, time = 0 min. (**C**) Appearance of the first DnaN-mCherry focus at the invasive pole of a *B. bacteriovorus* cell, indicating initiation of chromosome replication. (**D and E**) Replisome follows the newly replicated *oriC* region, indicating the progress of DNA replication. The appearance of a second DnaN-mCherry focus (reinitiation of chromosome replication), movement of the two replisomes toward opposite poles, and development of the third mNeonGreen-ParB focus in the middle of the filament. (**F and G**) Reinitiation of DNA replication from the chromosome located at the flagellar pole. Appearance of another ParB signals reflects the emergence of a newly replicated *oriC* region. (**H**) Termination of DNA replication. Photos represent merged bright-field and fluorescence (red and green) images. The *B. bacteriovorus* cell and the bdelloplast are marked by yellow and blue dotted lines, respectively. Scale bar = 1 μm. The full time-lapse is shown in Video S6.

**Figure 7.**
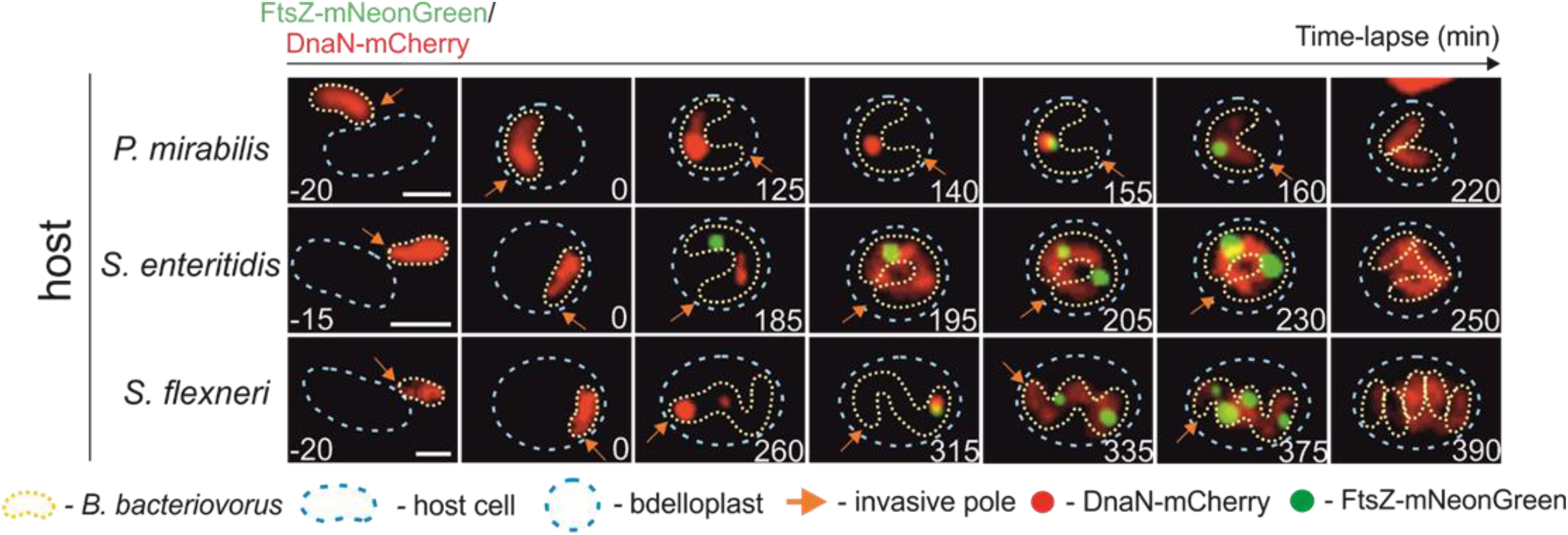
Binary and non-binary proliferation of *Bdellovibrio bacteriovorus* in hosts differing in size. (**A**) <u>Choreography of DNA replication initiation and modes of cell division.</u> Depending on the host cell’s size, *B. bacteriovorus* divides either by binary (in small host cells) or non-binary (in larger host cells) fission to produce two or more than two daughter cells, respectively. Regardless of the cell division mode, chromosome replication is always initiated at the invasive pole. In larger hosts where five or more progeny cells are formed, the last round(s) of chromosome replication may also be initiated at the cell pole opposite the invasive pole. (**B**) <u>Cell cycle of *B. bacteriovorus* dividing by binary fission, highlighting the dynamics of chromosome replication and cell division.</u> Chromosome replication is initiated at the invasive pole and one of the newly replicated *oriC* regions segregates toward the flagellated pole. During DNA replication, the replication forks split (one replisome follows the segregated *oriC* region), merge at the midcell and then disassemble. FtsZ ring assembly is initiated at the midcell before the termination of DNA replication, the filament subsequently divides into two progeny cells.

### Chromosome replication and segregation

Chromosome replication and segregation dynamics of *B. bacteriovorus* was studied in *P. mirabilis,* being the simplest model as only two daughter cells are produced (Fig. 1). Firstly, we intended to observe the localization of the replisome(s) during the reproductive phase. We therefore replace the native DnaN of *B. bacteriovorus* with a DnaN-mNeonGreen. The appearance and disappearance of DnaN-mNeonGreen foci indicate the assembly and disassembly, respectively, of replisome complexes and thus correspond to the initiation and termination of DNA replication (26). DNA synthesis started (i.e., a DnaN-mNeonGreen focus appeared) consistently at the invasive pole in bdelloplasts (n=100, 96%; Fig. 2C, Video S1) at 46±4 min after bdelloplast formation (Tab. 2). The second focus appeared at 31±4 min (n=100) after the first initiation of DNA replication, and subsequently migrated to the opposite pole (the pole that contained the flagellum before host attack, hereinafter called the ‘flagellar pole’). The pole-to-pole distance was covered in 11±2 min (n=100). Next, the replisomes relocated from the cell poles to a midcell position (Fig. 2, Video S1). Both replisomes remained at this position until the end of chromosome replication, which was indicated by diffusion of the DnaN-mNeonGreen fluorescent foci. It is worth noting that we never observed more than two foci during the reproductive phase in *P. mirabilis-derivative* bdelloplasts (n=100). Thus, the DnaN-mNeonGreen fluorescent foci presumably represent the two replication forks that, during DNA synthesis, transiently split at the invasive pole and then merge in the midcell position. The average duration of chromosome replication was 144 ±7 min (n=100).

We next investigated the dynamics of chromosome segregation in relation to DNA replication for *B. bacteriovorus* growing in *P. mirabilis*. For this purpose, we created allelic replacements of ParB and DnaN with fluorescent fusion proteins in *B. bacteriovorus* (see Table 1 and Table S1), which allowed us to simultaneously monitor both processes in a single cell. Notably, *B. bacteriovorus* mNeonGreen-ParB/DnaN-mCherry strain exhibited a proliferation-phase fluorescent pattern for DnaN-mCherry (Fig. 3, Video S2) consistent with that observed in *B. bacteriovorus* DnaN-mNeonGreen strain (Fig. 2, Video S1). In this system, DNA replication was followed by chromosome segregation (Fig. 3, Video S2): Shortly after the initiation of DNA synthesis (at 16±3 min and 69±4 min after bdelloplast formation, see Table 2), one of the newly replicated *oriC* regions visible as a ParB focus (ParB complexes colocalize with the *oriC* region; 18) started to migrate to the opposite cell pole, reaching its destination after 19±2 min (Table 2). In contrast to replisomes, the segrosomes did not relocate to midcell, but instead remained at the filament poles (Fig. 3, Video S2). Careful examination of dozens of proliferating *B. bacteriovorus* cells (n= 45; as an example, see Fig. 4, Video S3) revealed that the cell pole that was flagellar during the attack phase later became an invasive pole with a visible ParB complex. Thus, the two poles arising after division become the flagellar poles. The daughter cell which inherits the flagellar pole from the mother cell converts it into an invasive pole (Fig. 4, Video S3).

**Table 2.** Comparison of the average appearance times of replisome(s) and segrosome(s) in *B. bacteriovorus* cells during proliferation in different hosts.

| Average time*<br>[min]<br>HOST | 1<sup>st</sup> DnaN<br>focus | 2<sup>nd</sup> ParB<br>focus | Migration of<br>new *oriC* to<br>the opposite<br>pole | 2<sup>nd</sup> DnaN<br>focus | 3<sup>rd</sup> ParB<br>focus | 3<sup>rd</sup> DnaN<br>focus | 4<sup>th</sup> ParB<br>focus | Duration<br>of life cycle |
| --- | --- | --- | --- | --- | --- | --- | --- | --- |
| *P. mirabilis* | 46±4 | 69±4 | 19±2 | 79±6 | X | X | X | 252±7 |
| *S. enteritidis* | 46±5 | 65±5 | 18±2 | 119±7 | 138±8 | 163±9 | 182±10 | 288±7 |
| *S. flexneri* ** | 44±5 | 76±6 | 17±2 | 128±7 | 144±7 | 191±9 | 211±8 | 379±11 |

To understand how chromosome replication and segregation dynamics are affected by the production of multiple progeny cells we fed *B. bacteriovorus* on *S. enteritidis* (usually 3 or 4 daughter cells) and *S. flexneri* (4-8 daughter cells) (see Fig. 1). As expected, in these hosts, chromosome replication was initiated at the invasive cell pole of *B. bacteriovorus* (Figs. 5 and 6 and Fig. S5, Videos S4-S6), at 46±5 min after bdelloplast formation in *S. enteritidis* and at 44±5 min in *S. flexneri* consistent with previous experiments (Table 2). In *S. enteritidis* and *S. flexneri,* one of the newly replicated *B. bacteriovorus oriC* regions (i.e., an mNeonGreen-ParB focus) also migrated to the opposite cell pole, reaching its destination at 18±2 and 17±2 min, respectively, after *oriC* duplication (Videos S4-S6, Table 2). Meanwhile, the DnaN-mCherry focus started to move from the invasive pole and transiently followed the ParB focus, indicating the progression of mother chromosome replication (Fig. 5D, Fig. 6D, Fig. S5D, Videos S4-S6). The next steps in chromosome multiplication were related to the number of released daughter cells. When three progeny cells were produced during predation on *S. enteritidis*, DNA replication was reinitiated from the chromosome located at the invasive pole. The newly assembled replisomes (visible as a large DnaN-mCherry focus; see Fig. 5E, Video S4) moved away from the invasive pole (Fig. 5F-G, Video S4) and the first round of DNA replication was completed (Fig. 5G, Video S4). It was challenging to analyze the later stages of chromosome replication, because in most cases it was unclear whether the observed foci were from transiently splitting sister replication forks and/or from temporally overlapping replication rounds. When four daughter cells were formed in an *S. enteritidis* bdelloplast, we observed two asynchronous reinitiation events; in both cases, new replisomes were assembled at the invasive pole (see Fig. S5H-I, Video S5). Notably, the first reinitiation of DNA replication started (Fig. S5H) after the disappearance of the first DnaN-mCherry focus, which was located in the middle of the filament and originated from the first initiation.

In *S. flexneri* bdelloplasts, we observed up to five DnaN-mCherry foci, indicating that three (or more) pairs of replication forks were simultaneously active in a single filament. Unexpectedly, in an *S. flexneri* bdelloplast producing five (or more) progeny cells, at later stages of chromosome multiplication (150 min after bdelloplast formation; Fig. 6F, Video S6), we noticed the appearance of a DnaN-mCherry focus at the flagellar pole, suggesting that replication may also initiate at the opposite pole. Similar to the situation in *S. enteritidis* an observation of the later timepoints was not possible due to long, twisted filaments formation by the mother cells prior to cell separation.

In all bacteria, replication initiation is synchronized with the segregation of newly synthesized *oriC* region(s) (27). In all the hosts analyzed in our study, shortly after initiation of the first replication round, one of the newly synthesized *oriC* regions visible as a ParB focus (2^nd^ ParB focus) moved towards the flagellar pole (Figs. 3, 5, 6, Fig. S5, Table 2, Videos S2, S4-S6). In *S. enteritidis* or *S. flexneri,* similar to *E. coli* (18), the number of ParB foci gradually increased as new copies of the *oriC* region were synthesized (Table 2). The second round of chromosome segregation started at 74±5 min after the first round in *S. enteritidis,* and at 69±5 min in *S. flexneri* (n=100, data not shown). The newly assembled segrosome (3^rd^ ParB focus) also migrated towards the opposite pole but did not reach the end of the filament, which was already occupied by the 2^nd^ ParB complex; the 3^rd^ ParB focus remained at the midcell (Figs. 5, 6, Fig. S5, Videos S4-S6). After initiation of the third round of replication, the newly formed ParB focus (4^th^ ParB focus) moved away from the invasive pole to its final location between the 1^st^ and 3^rd^ ParB foci (Fig. 6F and S5J, Table 2). Careful examination confirmed that the number of ParB foci reflected the number of released daughter cells (n=300).

In conclusion, the choreography of *B. bacteriovorus* chromosome replication occurring in *S. flexneri* differs from those occurring in *P. mirabilis* and *S. enteritidis*: In *S. flexneri*, DNA replication may be reinitiated from the chromosome located at the flagellar pole.

### Septation

In all living cells, chromosome replication must be synchronized with cell division. Since chromosome replication is not followed by cell division in *B. bacteriovorus* except in cases of binary fission (see below), we decided to elucidate how chromosome replication is synchronized with filament septation during the growth of *B. bacteriovorus* in hosts that produce different numbers of daughter cells. For this purpose, we used a *B. bacteriovorus* strain producing DnaN-mCherry and FtsZ-mNeonGreen fusion proteins (Table 1). FtsZ filaments assemble into a ring (called the Z-ring) at the future site of the septum, and the appearance of a FtsZ focus is commonly regarded as an early marker of bacterial cell division (28). FtsZ from *B. bacteriovorus* is highly similar to other FtsZ homologs (53% identity with *E. coli* FtsZ) within the N-terminal domain, which contains conserved residues responsible for GTP binding, but possesses a unique C-terminal addition of ~160 amino acids (Fig. S6).

As expected, during predation in *P. mirabilis* only one FtsZ signal appeared before the *B. bacteriovorus* filament divided into two daughter cells. Recruitment of FtsZ-mNeonGreen to the division site started 165±4 min (n=100) after bdelloplast formation (Table 3, Fig. S7AG, Fig.7). The FtsZ focus was localized in the middle of the mother cell (Fig. 7, Fig. S7AB, Video S7), and was visible for 49±3 min (n=100, see Tab. 3). After FtsZ disassembly, the mother cells divided into two daughter cells. The duration of the *B. bacteriovorus* life cycle (from bdelloplast formation to progeny cell release) in *P. mirabilis* lasted approximately 4 hours (Table 2).

**Table 3.**
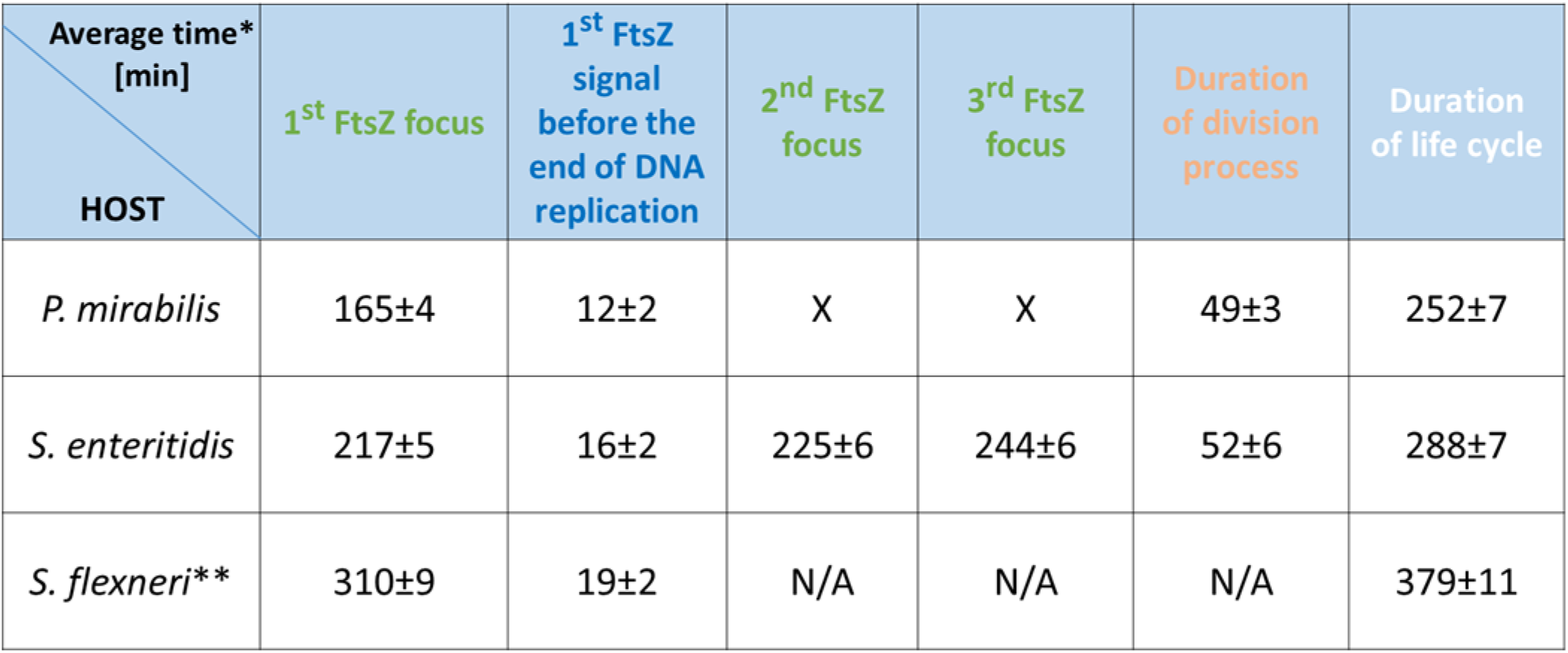
Comparison of the average times at which divisome(s) appeared in *B. bacteriovorus* cells during the septation process in different hosts.

| Average time*<br>[min]<br>HOST | 1<sup>st</sup> FtsZ focus | 1<sup>st</sup> FtsZ<br>signal<br>before the<br>end of DNA<br>replication | 2<sup>nd</sup> FtsZ<br>focus | 3<sup>rd</sup> FtsZ<br>focus | Duration<br>of division<br>process | Duration<br>of life cycle |
| --- | --- | --- | --- | --- | --- | --- |
| *P. mirabilis* | 165±4 | 12±2 | X | X | 49±3 | 252±7 |
| *S. enteritidis* | 217±5 | 16±2 | 225±6 | 244±6 | 52±6 | 288±7 |
| *S. flexneri* ** | 310±9 | 19±2 | N/A | N/A | N/A | 379±11 |

Investigation of the septation process during predation on *S. enteritidis* revealed that two or three FtsZ signals were usually visible inside a single filament (Fig. 7, Fig. S8, Video S8) and, after their disappearance, the filament septated into three (see Fig. 7 and Fig. S8) or four daughter cells, respectively. Surprisingly, in filaments growing in a host producing three or more sibling cells, the FtsZ foci appeared asynchronously (for details see Fig. 7, Figs. S8-9 and Table 3). For example, in *S. enteritidis* producing four daughter cells, the first FtsZ focus was visible 217±5 min after bdelloplast formation while the last one (i.e., the third one) appeared approximately half an hour later (244±6 min after bdelloplast formation) (Table 3). When three progeny cells were created, the first FtsZ focus was always localized closer to the flagellar pole (Fig. S8G). The FtsZ foci remained noticeable for 52±6 min and then underwent disassembly, and the filament then divided into four daughter cells.

In the case of *B. bacteriovorus* growing in *S. flexneri,* it was difficult to precisely analyze the septation process as the long, twisted filaments formed in this host prevented in depth microscopic analysis. At the beginning of the septation process, we were able to observe the sequential appearance of multiple FtsZ signals (up to 7) and, as in other hosts, the FtsZ rings never formed simultaneously (see for example Fig. S9, Video S9).

To investigate the septation process in relation to DNA replication we used TLFM analysis. The recruitment of FtsZ to the Z ring started before the end of DNA synthesis: In all hosts, the first FtsZ-mNeonGreen focus was visible approximately 10-20 minutes before the disassembly of the last replisome (Table 3, Fig. 7, Videos S7-S9), namely at 12±2 min in *P. mirabilis* (85%, n=100), 16±2 min in *S. enteritidis* (88%, n=100) and 19±2 min in *S. flexneri* (90%, n=100).

At the end of DNA synthesis, in binary dividing cells, the FtsZ-mNeonGreen focus consistently colocalized with DnaN-mCherry at the midcell position (n=100). This is simultaneously the junction point of the two replication forks (see the overlapped FtsZ-mNeonGreen and DnaN-mCherry signals on Fig. 7, Fig. S7, and Video S7). Interestingly, when three daughter cells were produced in an *S. enteritidis* bdelloplast, the first FtsZ ring never colocalized with the last visible replisome (i.e., the second and third DnaN focus that appeared, depending on the number of progeny cells). At the end of replication, the DnaN focus was located asymmetrically with respect to the filament length: It was closer to the invasive pole (94%, n=100). Meanwhile, the first FtsZ focus was visible near the flagellar pole (Fig. 7), where the first round of replication and segregation had already been completed (see Fig. 7, Fig. S8 and Table 3).

In *S. flexneri,* we were unable to precisely monitor the position of FtsZ rings in relation to the last visible replisome, especially in long filaments. However, we noticed that in a filament that would divide into five progeny cells, the first FtsZ-mNeonGreen focus frequently colocalized with the last visible DnaN-mCherry focus (78%, n=100, Fig. 7, Fig. S9G), which was positioned close to the flagellar pole (consistent with observations in *S. enteritidis*). We also observed that in *S. flexneri* producing five (or more) progeny cells, the initiation of replication may also take place at the flagellar pole (at later stages of chromosome multiplication; see Fig. 6F, 150 min after bdelloplast formation, Video S6).

Since we found that the formation of FtsZ rings in the predatory filament is an asynchronous process and the number of released *B. bacteriovorus* cells depends on the host/cell size, we decided to analyze the size of the progeny cells released from the analyzed pathogens in comparison to the model host, *E. coli*. Length measurement of mature *B. bacteriovorus* cells after bdelloplast lysis showed that there were statistical relevant differences (see Fig. S10): The average lengths of the generated *B. bacteriovorus* cells were 1.25±0.03 μm for *P. mirabilis,* 1.16±0.03 μm for *S. enteritidis,* 1.35±0.03 μm for *S. flexneri,* and 1.11±0.04 μm for *E. coli*. On average, the longest progeny cells were released from *S. flexneri* (usually 5-6 progeny cells), which had the largest cell size of the analyzed hosts. The daughter cells born from *P. mirabilis* (2 progeny cells) were longer than those released from *E. coli* (usually 3-4 progeny cells) and *S. enteritidis* (usually 3-4 progeny cells). There is therefore no direct correlation between the number of progeny cells and the size of the progeny cells.

Understanding the proliferation of *B. bacteriovorus* in different pathogens may also be helpful for its use to efficiently eliminate a given pathogen, i.e., by calculating the optimal ratio between the predator and the host. We developed a new application to simulate these infection dynamics using the number of predatory and prey cells, the average number of progeny cells released from a single host cell (Fig. 1) and the duration of the reproductive phase (Table 2). For the purposes of our simulation, we assume that all released progeny cells immediately find new hosts and that host cells do not replicate. The surviving number of host cells is calculated as n_host_i+1_ = n_host_i_ – n_bdellovibrio_i_ and the new number of *Bdellovibrio* cells is calculated as n_bdellovibrio_i+1_ = n_bdellovibrioi * n_progeny. This simulation, called BdelloSim, allowed us to calculate the time necessary to eliminate the entire host population given starting population sizes of 500000 and 100 cells for the host and *B. bacteriovorus* cells, respectively, and experimentally obtained data: for *P. mirabilis* 4000 min, for *S. enteritidis* 2610 min and for *S. flexneri* 2660 min (Fig. S2).

Taken together, the results of our TLFM analyses revealed the cell cycle parameters of *B. bacteriovorus* filaments growing in three different pathogens. Our work demonstrates that the mode of the *B. bacteriovorus* cell cycle depends on the host cell size: Filaments proliferating in small cells, such as *P. mirabilis*, divide by binary fission, while those growing in larger host cells undergo non-binary fission.

## Discussion

The use of predatory bacteria such as *B. bacteriovorus* is currently regarded as a promising strategy to kill antibiotic-resistant pathogens (7, 29, 30). Therefore, understanding the mode and dynamics of *B. bacteriovorus* proliferation in various dangerous pathogens may facilitate knowledge-based improvements in *B. bacteriovorus* strains for use as a “living antibiotic” in human and veterinary medicine, particularly to combat multidrug-resistant pathogens. Previously, *B. bacteriovorus* was believed to undergo a non-binary life cycle. In this study, we demonstrate for the first time that this predatory bacterium proliferates by either a binary or a non-binary mode of fission, depending on the host and its size (Fig. 8A). We also show that the choreography of DNA replication in this system exhibits unique features and is related to the mode of fission and/or number of progeny cells (Fig. 8A). Finally, we show that FtsZ rings are assembled asynchronously in non-binary proliferating cells of *B. bacteriovorus*.

**Figure 8.**
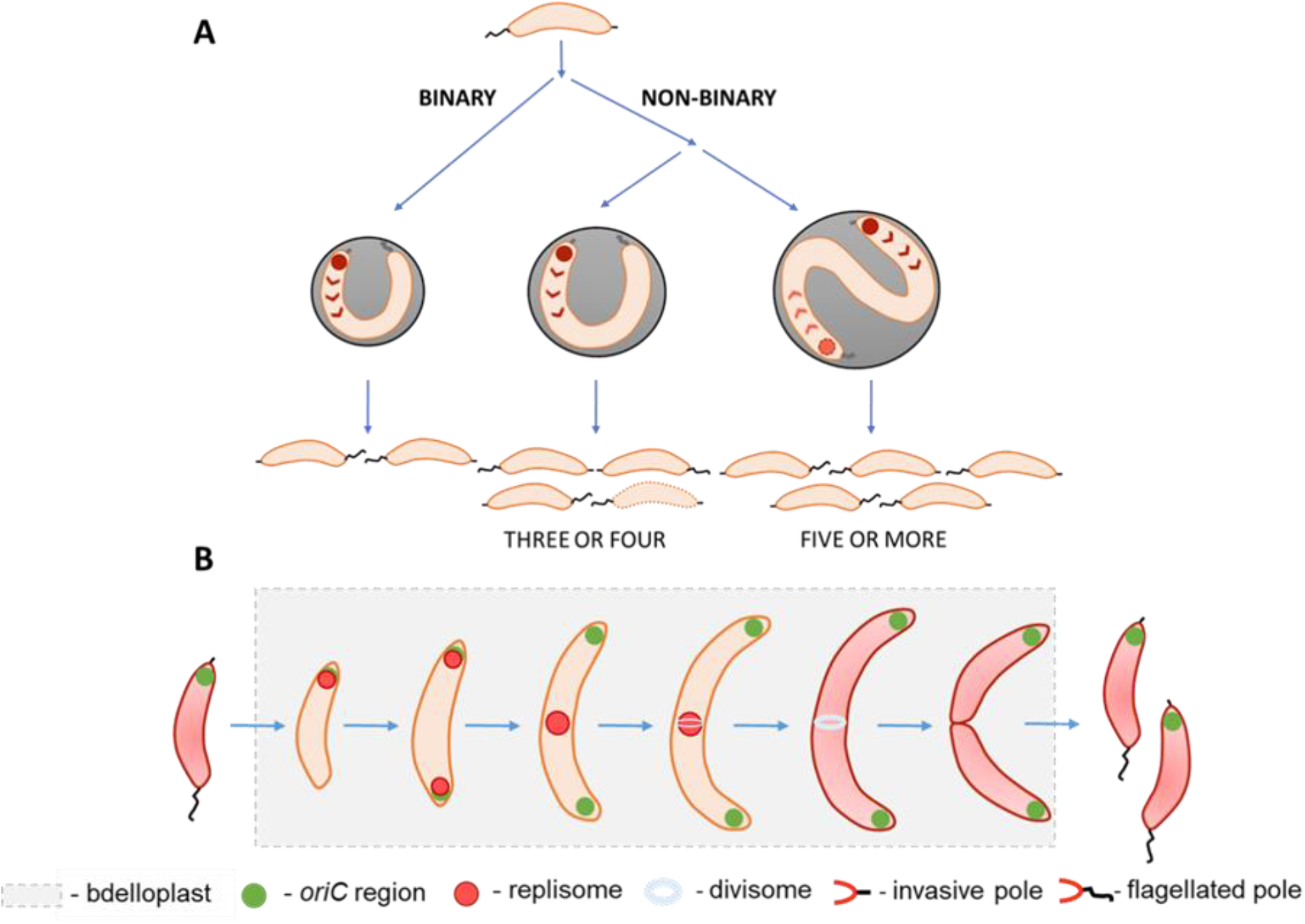
Binary and non-binary proliferation of *Bdellovibrio bacteriovorus* in hosts differing in size. (**A**) **Choreography of DNA replication initiation and modes of cell division**. *B. bacteriovorus* depending on a host size, divides either by binary (in small host cells) or by non-binary fission (in larger host cells) producing two or more daughter cells, respectively. The chromosome replication, regardless of the mode of cell division, is always initiated at the invasive pole. In larger hosts (when 5 or more progeny cells are formed), the last round(s) of chromosome replication may also be initiated at the cell pole, which is opposite to the invasive pole. (**B**) **Cell cycle of *B. bacteriovorus* dividing by binary fission, highlighting the dynamics of chromosome replication and cell division.** The chromosome replication is initiated at the invasive pole and then one of the newly replicated *oriC* regions is segregated toward the flagellated pole. During the progress of DNA replication, the replication forks split (one replisome follows the segregated *oriC* region) and merge at the midcell and then disassembled. The FtsZ ring assembly is initiated at the midcell before the termination of DNA replication and next the filament is divided into two progeny cells.

### *B. bacteriovorus* undergoes binary fission when growing in a small host cell

Our data reveal that, in small pathogenic bacteria such as *P. mirabilis* (cell length < 2 μm), only two predatory cells are formed (see Fig. 8AB). In this host, the FtsZ ring is assembled in the middle of the filament and then undergoes disassembly, whereupon the filament is divided into two daughter cells. Interestingly, the assembly of the FtsZ ring begins before the termination of DNA replication. Detailed microscopic analysis of *P. mirabilis* undergoing predation by *B. bacteriovorus* reveals that the old pole that had been flagellated becomes an invasive pole with a visible ParB complex. Simultaneous analysis of both replication and segregation processes confirmed this observation: During ongoing DNA replication, one of the newly replicated *oriC* regions (observed as a ParB focus) migrates towards the opposite pole (the one bearing a flagellum before the host attack). Further microscopic observations demonstrated that, after *B. bacteriovorus* division, the released daughter cell with the invasive pole arising from the flagellar pole of the mother cell attaches to the next cell host using this cell pole and proliferates within this host. Thus, after the division by binary fission, one daughter cell inherits the old invasive pole while the other inherits the flagellar pole of the mother cell, which must be extensively rebuilt to become an invasive pole. Consequently, flagella must be formed at new poles emerging from the last division (Fig. 8).

Collectively, our results demonstrate for the first time that *B. bacteriovorus* may undergo binary fission and *P. mirabilis* provides a simple and ideal model host for studying the life cycle of this predatory bacterium.

### Unique choreography of chromosome replication in binary-dividing *B. bacteriovorus*

As in other hosts, including *E. coli* (17, 18), chromosome replication in *B. bacteriovorus* undergoing predation on *P. mirabilis* starts at the invasive pole. Then, shortly after the initiation of DNA synthesis, the replisomes split; one migrates towards the flagellar pole while the other remains at the invasive pole. Before the end of DNA synthesis, the two replisomes move towards each other to meet at the cell center, where assembly of the FtsZ ring has begun. Thus, at the beginning of chromosome replication, the choreography of this process in *B. bacteriovorus* resembles that found in other asymmetrically dividing bacteria, such as *C. crescentus* (31) and *V. cholerae* (chromosome I) (32), where the replisomes are also assembled at a cell pole. Contrary to chromosome replication in *E. coli* (27, 33) and *B. subtilis* (34), where replication starts at the midcell. However, the spatiotemporal choreography of further steps of *B. bacteriovorus* chromosome replication differs from those described in other asymmetrically dividing bacteria. For example, in *C. crescentus*, the replisomes migrate together from the cell pole to the midcell position, where replication is terminated (31).

The assembly of replisomes at the invasive pole of *B. bacteriovorus* (17, 18) suggests that, as in other asymmetric bacterial cells, the *oriC* region must be anchored to the cell pole via (a) specific polar protein(s). To date, such polar proteins, also called “hub” proteins, have been characterized in *C. crescentus* (PopZ) (35), *V. cholerae* (HubB) (36) and *Streptomyces* (DivIVA) (37, 38). The *B. bacteriovorus* genome lacks homologs of PopZ or HubP, but does encode a version of DivIVA (39). It was recently speculated that DivIVA together with a recently identified homolog of the filament-forming protein, bactofilin (39), may be involved in maintaining the polarity of this predatory bacterium. However, the precise role of these proteins in anchoring the *B. bacteriovorus oriC* region at the invasive pole remains to be elucidated. We cannot exclude the possibility that other proteins function to anchor the chromosome at the invasive pole.

In sum, we herein present for the first time the choreography of chromosome replication and cell division during the binary cell cycle of *B. bacteriovorus* (see Fig. 8B).

### The FtsZ rings are asynchronously assembled across the elongated filament during non-binary fission

In larger hosts (cell length > 2 μm), the filament of *B. bacteriovorus* undergoes non-binary division and odd or even numbers of cells are produced. As expected, the number of released progeny cells strictly depends on the number of FtsZ rings that are formed before filament division (Fig. 7). Although the septation process in *B. bacteriovorus* was previously shown to be synchronous (19), our study revealed that FtsZ is recruited asynchronously to the sites of division, with the first FtsZ signal appearing before the end of DNA synthesis (Fig. 7). Further FtsZ signals appear sequentially. For example, in a filament that will divide into four nascent cells, the third FtsZ signal is assembled ~ 30 minutes after the appearance of the first FtsZ signal (Table 3). In *Streptomyces,* which is a Gram-positive filamentous bacterium, the multigenomic filaments also undergo synchronous division to form chains of unigenomic exospores. However, in contrast to *B. bacteriovorus*, the FtsZ rings are synchronously assembled in this case (3, 40) and this occurs after the cessation of DNA replication (41). Thus, we speculate that in *B. bacteriovorus*, yet undiscovered replication checkpoints act to coordinate chromosome replication with cell cycle progression, such as by tuning FtsZ ring placement to prevent the formation of daughter cells with a missing/guillotined chromosome or an additional copy of the chromosome.

### In non-binary dividing long filaments, the initiation of chromosome replication may also occur at the cell pole opposite the invasive pole

Similar to the results obtained by Laloux’s group (18), we observed that DnaN foci usually appeared sequentially at the invasive pole, suggesting that there is asynchronous firing of replication initiation from the chromosome attached to this pole (the newly appeared fluorescently tagged DnaN foci colocalized with the ParB attached to the invasive pole; Figs. 5, 6, Fig. S5). However, it proved hard to follow the further progression of chromosome multiplication (especially in filaments that would divide into several or more nascent cells) because it was difficult to distinguish between transiently splitting sister replication forks and temporally overlapping replication rounds. In *Streptomyces* vegetative hyphae, DNA replication is also (re)initiated from the chromosome that is anchored at the filament tip, and one of the newly replicated *oriC* regions bound by ParB protein is unidirectionally segregated (4). Thus, in both *B. bacteriovorus* and *Streptomyces*, after *oriC* duplication, ParB complexes must be reestablished at both daughter *oriC*s at the cell pole; thereafter, one of them is captured by apically localized hub proteins and becomes abandoned by the segregation machinery.

Surprisingly, during the proliferation of *B. bacteriovorus* in *S. flexneri*, we noticed that the reinitiation of chromosome replication may also occur at the flagellar pole (Fig. 6, Video S6).

This phenomenon was observed only in the very late stages of chromosome multiplication, at ~ 150 minutes after the first initiation of chromosome replication. We speculate that in the longer filaments (those that will divide into five or more progeny cells), the flagellar pole is converted to an invasive pole while chromosome replication is still in progress. Therefore, the last round(s) of DNA replication may also be initiated from the chromosome that is already attached to the newly formed invasive pole. However, further work is needed to elucidate the mechanism(s) underlying the molecular switch that activates replication initiation from the chromosome where it attaches to the old invasive pole and/or the pole that has been converted from flagellar to invasive.

In sum, our findings demonstrate that in a non-binary dividing filament of *B. bacteriovorus*, chromosome replication may be initiated not only from the invasive pole, but also from the opposite pole, which has presumably been converted to an invasive pole.

### Conclusions

We herein set the scene for *B. bacteriovorus* to become a novel model that enables the study of both binary and non-binary cell division. Our findings provide new insight into the life cycle of a predatory bacterium that exhibits bimodal fission, and demonstrate that the mode of division depends on the size of the prey bacterium inside which *B. bacteriovorus* grows. Further studies on the *B. bacteriovorus* life cycle should aim to shed light on the regulatory mechanisms responsible for switching between the binary and non-binary modes of the cell cycle. Moreover, understanding the proliferation of *B. bacteriovorus* in different pathogens may facilitate its use to efficiently eliminate a given pathogen (i.e., by calculating the optimal ratio between the predator and the host).

## Materials and methods

### Bacterial strains and culture conditions

The culture conditions and construction of bacterial strains were as described in Makowski *et al*. (17). Briefly, the plasmids used to construct *B. bacteriovorus* HD100 strains were propagated in *E. coli* DH5α grown in LB broth or on LB agar plates (supplemented with 50 μg/mL kanamycin) and then transformed into *E. coli* S17-1. The *E. coli* strains used as prey for *B. bacteriovorus* were liquid cultured in YT medium (0.8% Bacto tryptone, 0.5% yeast extract, 0.5% sodium chloride, pH 7.5) with (S17-1 pZMR100) or without (S17-1, ML35) kanamycin (50 μg/mL), at 37°C with shaking (180 rpm). *B. bacteriovorus* cells were grown by predation on *E. coli* (ML35, S17-1 or S17-1 pZMR100) or pathogenic hosts (*P. mirabilis, S. enteritidis* or *S. flexneri)* in Ca-HEPES buffer (25 mM HEPES, 2 mM calcium chloride, pH 7.6). All pathogenic bacteria were grown in LB broth at 37°C with shaking (180 rpm) or on LB agar plates at 37°C. Details regarding the strains, plasmids and oligonucleotides used in this study are presented in Tables S1 and S2.

### Predatory killing curves

*B. bacteriovorus* strain HD100 (wild-type) was propagated in *P. mirabilis, S. enteritidis* or *S. flexneri*. An overnight culture of host cells was spun down at 5000 rpm for 10 min at 20°C, and the cells were washed with Ca-HEPES buffer and back diluted to OD_600_ = 1.0 with Ca-HEPES buffer. The cultures of *B. bacteriovorus* cells were filtered through 0.45 μm filters and 20 μL of filtered predator cells were mixed with 280 μL of host suspension (17). Lysis curves were analyzed using Bioscreen C (Automated Growth Curves Analysis System, Growth Curves, USA). The decrease of optical density (OD_600_) at 30°C was measured at 20-minute intervals for 42 hours. Experiments were done in three independent biological replicates. The Weibull four-parameters model was fitted to the data using the drc R package to calculate EKT_50_ values and their standard errors (22).

### DNA manipulation and construction of *B. bacteriovorus* HD100 strains

DNA manipulations in *E. coli* were carried out using standard molecular methods (23). Oligonucleotides were synthesized by Sigma-Aldrich. Reagents and enzymes were supplied by Thermo Scientific, Sigma-Aldrich and NEB. The following *B. bacteriovorus* HD100 strains were constructed: DnaN-mNeonGreen, DnaN-mCherry, mNeonGreen-ParB/DnaN-mCherry and FtsZ-mNeonGreen/DnaN-mCherry. In them, the replisomes were labeled green or red by DnaN fusion or the segrosomes and divisomes were labeled green by ParB and FtsZ fusions, respectively (Table S1).

The coding sequences of the relevant *B. bacteriovorus* genes *(dnaN, parB* or *ftsZ)* or reporter genes (*mNeonGreen* or *mCherry*) were amplified (all primers are listed in Table S2) using chromosomal DNA isolated from *B. bacteriovorus* HD100 or plasmids (pAKF220-mNeonGreen or p2Nil-lsr2-mCherry), respectively, as the templates. Gibson assembly was used to clone the PCR products into the pK18mobsacB plasmid. The obtained constructs (pK18dnaN-mNeonGreen, pK18dnaN-mCherry, pK18mNeonGreen-parB and pK18ftsZ-mNeonGreen) were transformed into *E. coli* S17-1 and conjugated to *B. bacteriovorus* strain HD100 as described previously (17). Scarless allelic replacements of the wild-type chromosomal *dnaN, parB* and *ftsZ* genes by the gene fusions were performed using two-step homologous recombination with suicide vector pK18mobsacB, as described by Jurkevitch *et al*. (24). Double-crossover (DCO) events were performed on single-crossover (SCO) strains of *B. bacteriovorus*. SCO strains were grown for 2 days without antibiotics in Ca-HEPES and then for an additional 2 days in Ca-HEPES with 5% sucrose. The obtained mutants were diluted and grown on two-layer YPSC agar plates (1% Broadbeam peptone, 1% yeast extract, 0.5% sodium acetate, 0.25% magnesium sulfate, [pH=7.6]). The final concentrations of agarose in the YPSC plates were 1% in the bottom layer and 0.6% in the top layer. The plates were incubated at 30°C until plaques appeared. DCO strains were searched by plaque PCR screening. The obtained *B. bacteriovorus* strains harbored gene fusions under the control of the endogenous promoters. Proper construction of all fluorescent strains was verified by DNA sequencing and fluorescent microscopic observations.

### Microscopic analysis

Agarose gel (1%) in Ca-HEPES buffer was poured onto a basic slide and allowed to solidify. Lysate (10 μL) of fresh prey (wild-type *E. coli, P. mirabilis, S. enteritidis* or *S. flexneri)* was added in a layer of agarose gel and the slide was sealed with a coverslip. Images were taken in phase-contrast mode using a Leica DM6 B microscope equipped with a 100x/1.32 OIL PH3 objective. Captured images were analyzed using the ImageJ Fiji suite (http://fiji.sc/Fiji).

### Time-lapse fluorescence microscopy

Time-lapse fluorescence microscopy (TLFM) was performed on B04A plates with an ONIX flow control system (Merck-Millipore) (24) or on an agarose pad (25). For the ONIX system, an overnight stationary-phase culture of host cells was diluted 25 times and loaded into the flow chamber. Next, *B. bacteriovorus* cells were injected to the flow chamber (pressure, 4 lb/in^2^) in Ca-HEPES buffer at 30°C for 2 min or until predatory cells were visible in the field of view. Agarose pads were prepared as described previously (17, 25) using *P. mirabilis* as a prey cell. In both methods, images were recorded every 5 min using a Delta Vision Elite inverted microscope equipped with an Olympus 100x/1.40 objective and a Cool SNAP HQ2-ICX285 camera. DnaN-mCherry was visualized by detection of mCherry (EX575/25; EM625/45) with an 80-ms exposure time and 50% intensity. Proteins fused with mNeonGreen were visualized by detection of green fluorescent protein (GFP; EX475/28; EM525/48) with a 50-ms exposure time and 32% intensity. Bright-field images were taken with a 30-ms exposure time and 10% intensity. The captured images were analyzed using the ImageJ Fiji suite (http://fiji.sc/Fiji). TLFM experiments were done in three independent biological replicates.

### Statistical analysis and *B. bacteriovorus* growth simulation

All obtained values and time points were calculated from at least hundred *B. bacteriovorus* cells using using three independent time-lapse microscopy experiments. For all measured values, the 95% confidence interval (CI) was calculated, assuming a normal distribution of results. Simulations of *B. bacteriovorus* population growth were done using the R language based on the experimental data concerning the life cycle length and number of progeny cells. The code is available at https://github.com/astrzalka/BdelloSim and the online version of the application, which allows for simultaneous analysis of three different hosts, is available at http://microbesinwroclaw.biotech.uni.wroc.pl:3838/BdelloSim/.

## Acknowledgments

This work was financed by a National Science Centre grant Opus (2018/29/B/NZ6/00539). The authors are grateful to Bernhard Kepplinger and Dagmara Jakimowicz for critically reviewing the manuscript.

