## Supplementary materials for "Binary or non-binary fission? Reproductive mode of a predatory bacterium depends on prey size"

**Table S1** Bacterial species, strains and plasmids used in this study.

| Bacterial species, strain and plasmid | Description/sequence | Source |
|---|---|---|
| **Bacterial species** | | |
| ***E.coli*** | | |
| S17-1 | thi pro hsdR$^-$ hsdM$^+$ recA; harboring plasmid RP4-Tc::Mu-Kn::Tn7, used as donor for conjugation of plasmids into *Bdellovibrio* | (43) |
| S17-1 pZMR100 | S17-1 strain containing pZMR100 plasmid to confer Kan$^r$ ; used as Kan$^r$ prey for *Bdellovibrio* | (42) |
| ML35 | Strain routinely used as prey for *B. bacteriovorus* | ATCC43827 |
| *Proteus mirabilis* | Non-pathogenic strain used as prey for *Bdellovibrio* | PCM1098 |
| *Salmonella enteritidis* | Non-pathogenic strain used as prey for *Bdellovibrio* | PCM2550 |
| *Shigella flexneri* | Non-pathogenic strain used as prey for *Bdellovibrio* | PCM1936 |
| ***B. bacteriovorus* strains** | | |
| HD100 | Wild-type strain | DSMZ50701 |
| HD100DnaN-mCherry | HD100 *dnaN::dnaN-mCherry* | This study |
| HD100DnaN-mNeon | HD100 *dnaN::dnaN-mNeonGreen* | This study |
| HD100mNeon-ParB/DnaN-mCherry | HD100 *parB::mNeonGreen-parB dnaN::dnaN-mCherry* | This study |
| HD100FtsZ-mNeon/DnaN-mCherry | HD100 *ftsZ::ftsZ-mNeonGreen-ftsZ dnaN::dnaN-mCherry* | This study |
| **Plasmids** | | |
| p2Nil-*lsr2-mCherry* | Plasmid carrying mCherry coding sequence; Kan$^R$ | Marta Kołodziej |
| pAKF220 | Plasmid carrying mNeonGreen coding sequence; Amp$^R$ | (17) |
| pK18*mobsacB* | Suicide vector used for conjugation and recombination into *Bdellovibrio* genome; Kan$^R$ | (44) |
| pK18_dnaN_mCherry | Derivative of pK18mobsacB containing fusion gene *dnaN-mCherry*; Kan$^R$ | This study |
| pK18_dnaN_mNeon | Derivative of pK18mobsacB containing fusion gene *dnaN-mNeonGreen*; Kan$^R$ | This study |
| pK18_mNeon_ParB | Derivative of pK18mobsacB containing fusion gene *mNeonGreen-parB*; Kan$^R$ | This study |
| pK18_ftsZ_mNeon | Derivative of pK18mobsacB containing fusion gene *ftsZ-mNeonGreen*; Kan$^R$ | This study |

ATCC - American Type Culture Collection

PCM – Polish Collection of Microorganisms

DSMZ - Deutsche Sammlung von Mikroorganismen und Zellkulturen

**Table S2** Sequences of primers used in this study.

| ***B. bacteriovorus* strains** | **Name of primer** | **Sequence of primer (5 '→ 3')** |
|---|---|---|
| HD100DnaN-mCherry | pK18_DnaN_fw | ACCTCCGGGTACCGAGCTCGATGAAATTAGAGATTGATAAGCGAGATCTGTTAAGT |
| | linker_DnaN_rv | CCGCCGAGCCGATTCTCATTGGCATCACAACGCAG |
| | linker+mCherry_fw | GGCTCGGCGGGCTCGGCGGCGGGCTCGGGCGAGTTCATGGTGAGCAAGGGCGAGG |
| | recF-mCherry_rv | CGAAAATCATTACTTGTACAGCTCGTCCATGCC |
| | dnaN_linker_fw | AATGAGAATCGGCTCGGCGGGCT |
| | mCherry_RecF_fw | TGTACAAGTAATGATTTTCGAAAGACTGCGTCTGG |
| | pK18_RecF_rv | CAGCTATGACCATGATTACGCTACTCAAGGATTTGGCCATCCTTGA |
| | DnaN_upstream_fw | ATGGACTTAGGGGATTTGTTAGG |
| | FP_sek_rv2 | TACCCTGTCCGTTCTCTCCG |
| | DnaN_DCO_fw | ACTGTGGAAATAGCTGTGGAAGG |
| | RecF_DCO_rv | TACGAGCAACTCATCCACCAGC |
| HD100DnaN-mNeonGreen | pK18_DnaN_fw | ACCTCCGGGTACCGAGCTCGATGAAATTAGAGATTGATAAGCGAGATCTGTTAAGT |
| | linker_DnaN_rv | CCGCCGAGCCGATTCTCATTGGCATCACAACGCAG |
| | linker+mNeon_fw | GGCTCGGCGGGCTCGGCGGCGGGCTCGGGCGAGTTCATGGTTTCGAAAGGAGAGGAGG |
| | RecF_mNeon_rv | CGAAAATCATTTACTTGTACAGCTCGTCCATGCC |
| | dnaN_linker_fw | AATGAGAATCGGCTCGGCGGGCT |
| | mNeon_RecF_fw | CTATAAGTGAATGATTTTCGAAAGACTGCGTCTGG |
| | pK18_RecF_rv | CAGCTATGACCATGATTACGCTACTCAAGGATTTGGCCATCCTTGA |
| | DnaN_upstream_fw | ATGGACTTAGGGGATTTGTTAGG |
| | FP_sek_rv2 | TACCCTGTCCGTTCTCTCCG |
| | DnaN_DCO_fw | ACTGTGGAAATAGCTGTGGAAGG |
| | RecF_DCO_rv | TACGAGCAACTCATCCACCAGC |
| HD100mNeonGreen-ParB | pK18_parB_fw | GTCGACTCTAGAGGATCCCCTTACTGCCATCCTTCTTTAAGCCTATCTACC |
| | linker_parB_rv | GGGCGAGTTCATGTCTGATATTGCTGTAGAATCCTCAAACAAG |
| | mNeon_linker_rv | GAACTCGCCCGAGCCCGCCGCCGAGCCCGCCGAGCCCTTATAGAGTTCATCCATACC |
| | parA_mNeon_rv | GGGAATTTAAATGGTTTCGAAAGGAGAGGAGGATAATATG |
| | parB_linker_fw | TATCAGACATGAACTCGCCCGAGCCC |
| | mNeon_parA_fw | TCGAAACCATTTAAATTCCCCTTCTTATGCCATTTGTTCTG |
| | pK18_parA_rv | CGAATTCGAGCTCGGTACCCTTGAATGTTCACGCAAAAAGGAGGACT |
| | parB_zew_fw | TCTGAACCGTCCCCTCGAAGC |
| | parA_wew_SCO_rv | AATCCATCTTCGAGTACGACAGC |
| | parB_wew_fw | CGTACGACGAGCCACGATGG |
| | parA_wew_rv | ACCAGTATGACTTCGTGATCATCG |
| HD100FtsZ-mNeonGreen | linker_FtsZ_fw | GCCCGCCGAGCCTTCTTTATTCAGATCGAATCCTTGTTTCTTGCG |
| | pK18_FtsZ-rv | TACGAATTCGAGCTCGGTACCCGGGATGTTTGAGTTGGAAGAAAATATCAATATCGGTG |
| | IpxH_mNeon_fw | GGCTCGGCGGGCTCGGCGGCGGGCTCGGGCGAGTTCATGGTTTCGAAAGGAGAGGAGG |
| | linker_mNeon_fw | GGCTCGGCGGGCTCGGCGGCGGGCTCGGGCGAGTTCATGGTTTCGAAAGGAGAGGAGG |
| | IpxH_mNeon_fw | ACCAGGCTTCCACTACTTATAGAGTTCATCCATACCCATCACG |
| | FtsZ_linker_rv | TCTGAATAAAGAAGGCTCGGCGGGCT |
| | pK18_IpxH_fw | CCAGTGCCAAGCTTGCATGCCTGCATCATAAATCCTCCTCAGAAGGCAGA |
| | mNeon_IpxH_rv | ACTCTATAAGTAGTGGAAGCCTGGTTCATATCCGACAT |
| | FtsZ_downstream_SCO_fw | ATAGGTGCAGGTGTAAGAACGC |
| | FP_sek_rv2 | ATGGCTCGTGAAGTCCTGAGC |
| | FtsZ_downstream_fw, | TTCCCATTGCGCTCTTCAGCG |
| | FtsZ_DCO_upstream_rv | ATCGCTGGACGGTATCACC |

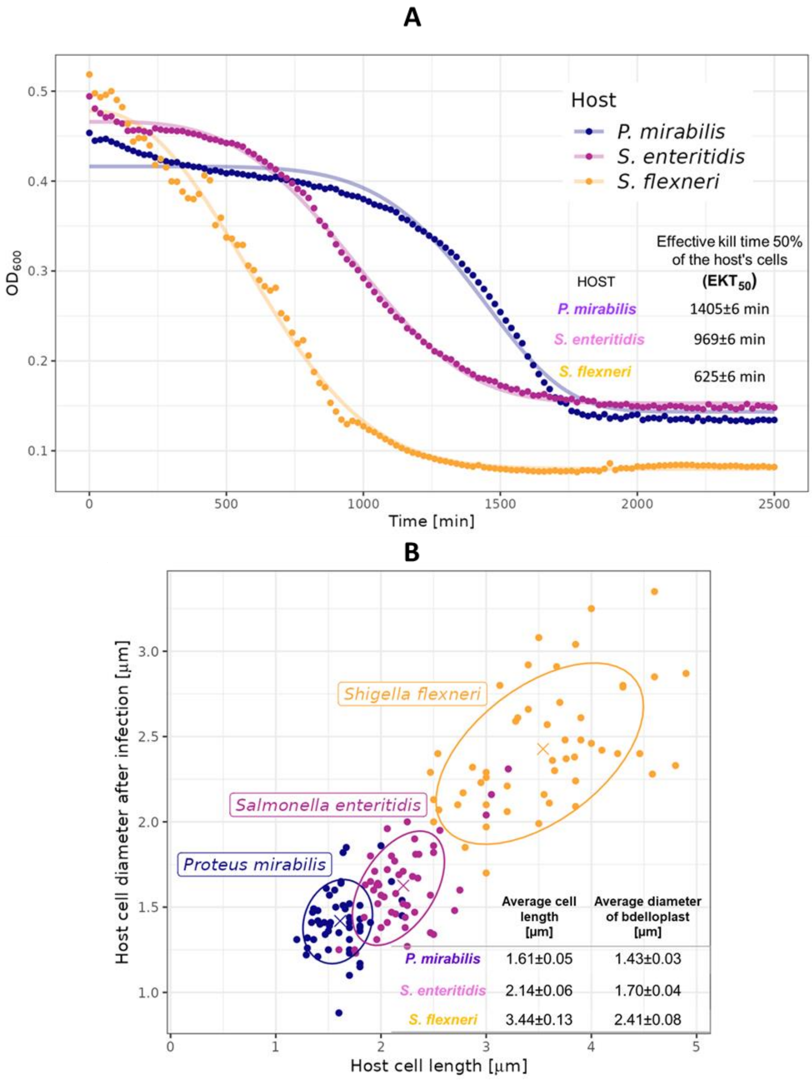


**Fig. S1. Basic features of *B. bacteriovorus* proliferation in different hosts.**

(**A**) Kill curves of *B. bacteriovorus* during predation on three different hosts: *P. mirabilis*, *S. enteritidis* and *S. flexneri*. Solid lines show the model fitted using a four-parameter Weibull function. (**B**) Correlation between the host cell length and the diameter of the bdelloplast formed after *B. bacteriovorus* enters the prey (n=100 for each host).

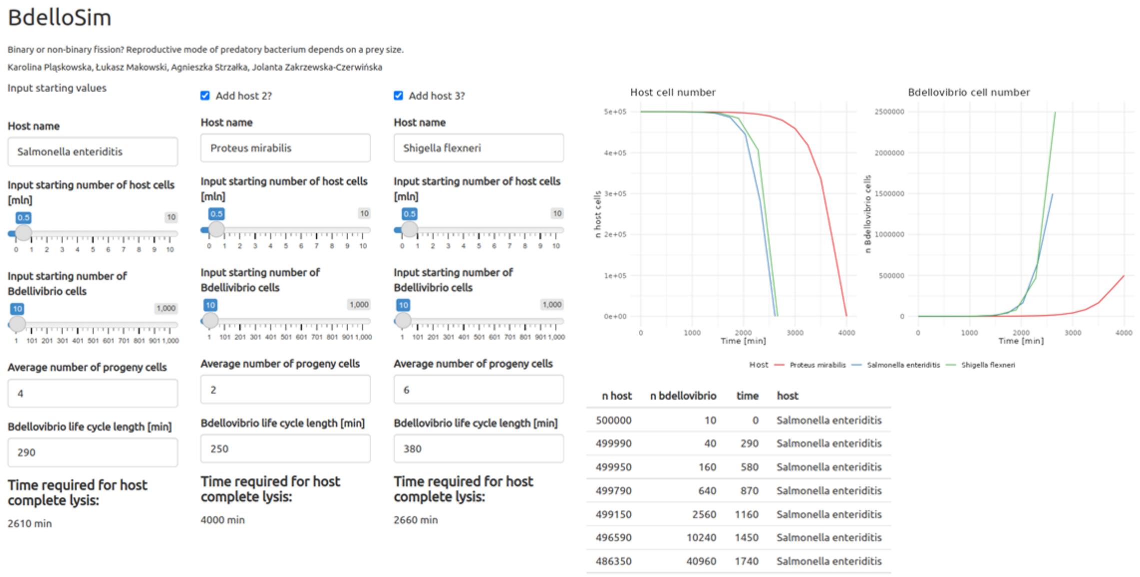


**Fig. S2. Screenshot from an application that simulates *B. bacteriovorus* population growth during proliferating in different cell hosts**. Simulations based on the experimental data concerning life cycle length and the number of progeny cells released from bdelloplast. The online version of the application is available at http://microbesinwroclaw.biotech.uni.wroc.pl:3838/BdelloSim/

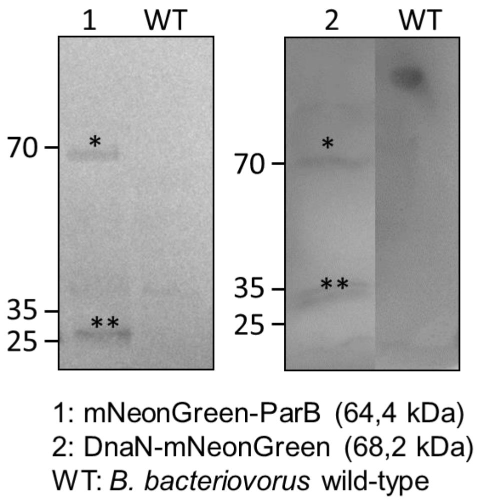


**Fig. S3.** Western blots of whole-cell protein extracts from *B. bacteriovorus* were probed with mNeonGreen antibody to confirm proper protein production. Line 1 and 2 are for mNeonGreen-ParB and DnaN-mNeonGreen, respectively. One star reflected detected proteins with fusion and double star reflected only fluorescent protein mNeonGreen (26,6 kDa). B. bacteriovorus HD100 (wild-type) was used as a negative control.

Western blot analysis

50 mL of overnight culture of *B. bacteriovorus* cells was spun down at 6000 rpm for 20 min at 20°C and resuspended in 5 mL fresh Ca-HEPES buffer. An overnight culture of *E. coli* ML35 cells was spun down at 5000 rpm for 10 min at 20°C, and the cells were washed and back diluted to $OD_{600}$ = 1.0 with Ca-HEPES buffer. The concentrated culture of *B. bacteriovorus* mixed with 4 mL of diluted *E. coli* cells and added 4 ml of Ca-HEPES buffer to final volume 12 mL. After 180 min of incubation, the culture was spun down and resuspended in 300 µL of Ca-HEPES buffer with protease inhibitor cocktail (Pierce Protease Inhibitor Tablets, Thermo Scientific). Cell lysate proteins (10 µg in total) were separated in a 10% denaturing polyacrylamide gel before being transferred to a nitrocellulose membrane. The protein was subsequently detected using primary mouse monoclonal anti-mNeonGreen antibodies and secondary goat anti-mouse IgG- antibodies conjugated with horseradich peroxidase. Signal from antibody binding was visualized by detecting chemiluminescence, which was imaged with Biorad Universal Hood II Gel Doc System.

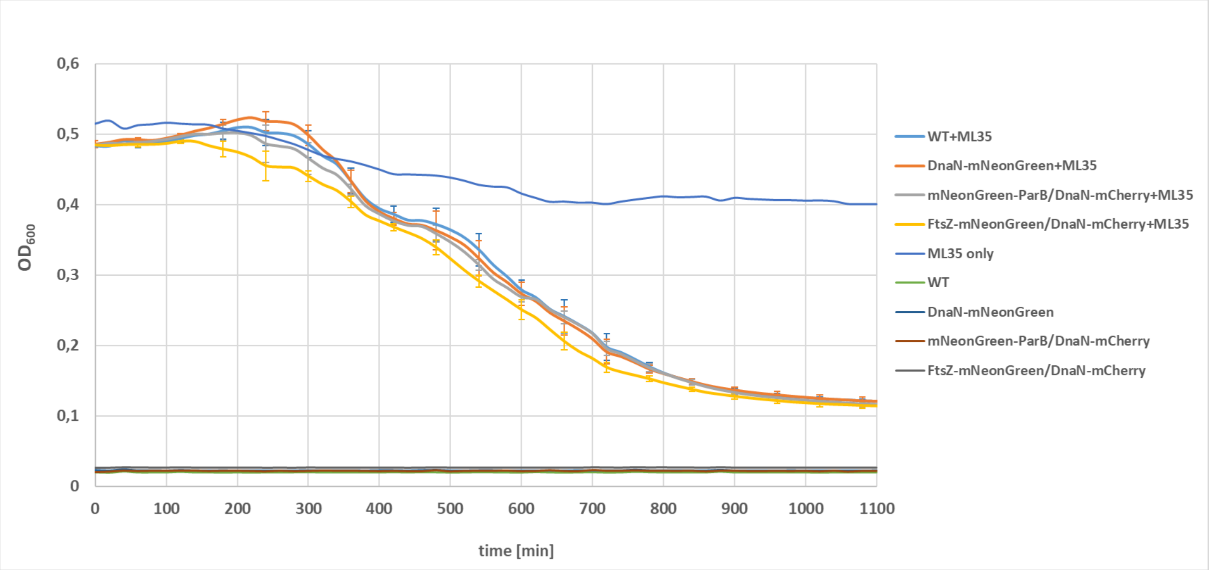


**Fig. S4. Kill curves of *B. bacteriovorus* strains**. Predation of *B. bacteriovorus* strains DnaN-mNeonGreen, mNeonGreen-ParB/DnaN-mCherry, FtsZ-mNeonGreen/DnaN-mCherry and wild-type on *E. coli* ML35. Obtained kill curves are similar, which indicates that genetic manipulations in *B. bacteriovorus* strains did not affect the effectiveness of lysis of the prey.

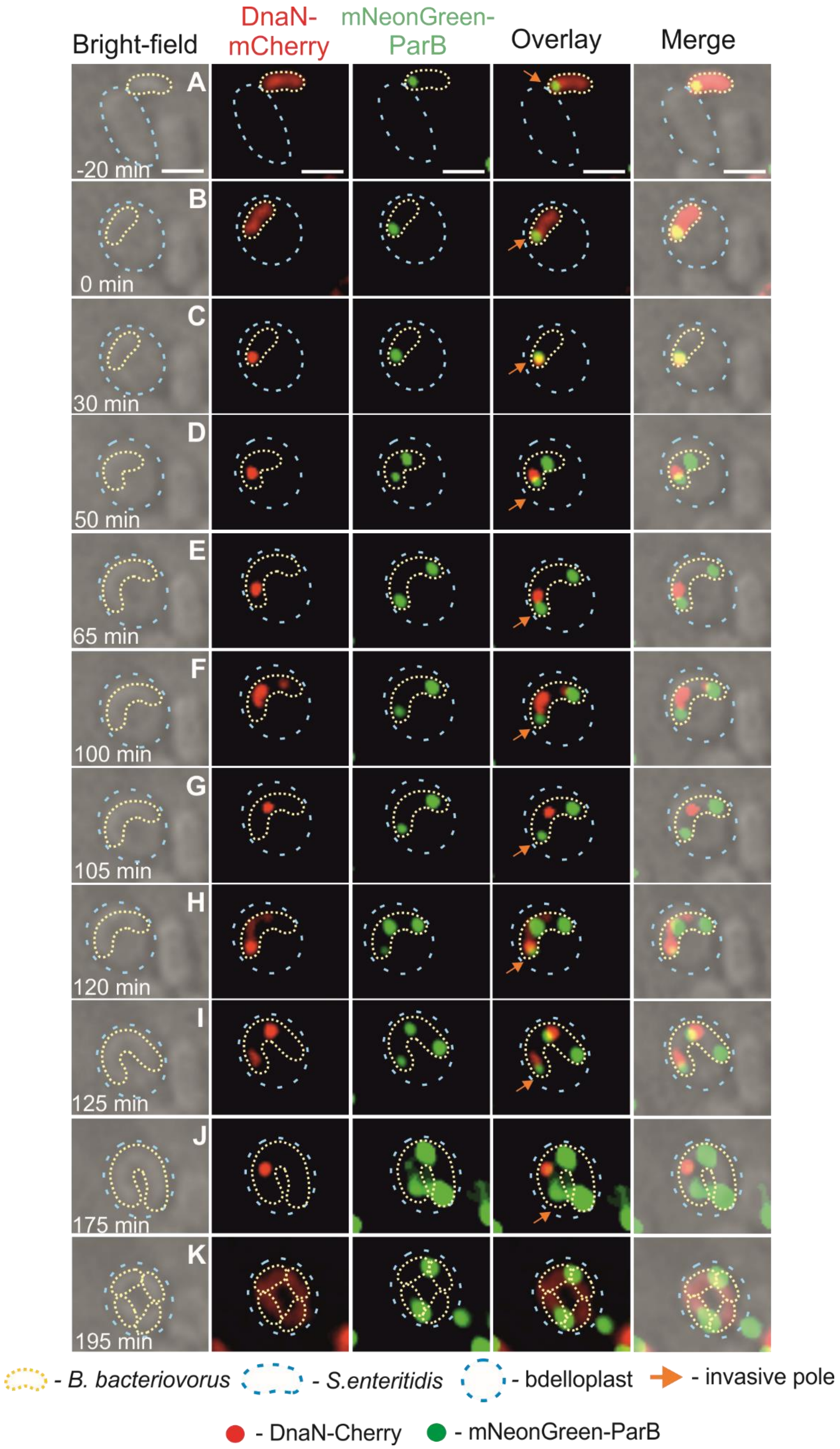


**Fig. S5. Dynamics of chromosome replication and *oriC* segregation in a *B. bacteriovorus* cell during proliferation in *S. enteritidis* – formation of four daughter cells.** (**A**) Attachment of *B. bacteriovorus* to a *S. enteritidis* cell. (**B**) Bdelloplast formation, time = 0 min. (**C**) Appearance of the first DnaN-mCherry signal at the invasive pole - initiation of chromosome replication. (**D-G**) Replisome movement behind the ParB focus, indicating the progress in DNA multiplication. The appearance of second DnaN-mCherry foci, which presumably reflect splitting two replication forks and next merge in the middle of filament. (**H-I**) Two reinitiations of DNA replication from the mother chromosome located at the invasive pole and migration of both replisomes toward the mid-cell position. At the same time appearance of the third signal of ParB, which was located in the middle of the filament. (**J**) The disappearance of one replisome located nearest the old flagellar pole - termination of one chromosome replication and appearance of fourth mNeonGreen-ParB foci, which was located between signals at the invasive pole and in the middle of predator cell, respectively. (**K**) Termination of chromosomes replication and formation of four daughter cells.

Photos represent merged bright-field and fluorescence (red and green) images. The *B. bacteriovorus* cell and the bdelloplast are marked by yellow and blue dotted lines, respectively. Scale bar = 1 µm. The full time-lapse is shown in Video S5.

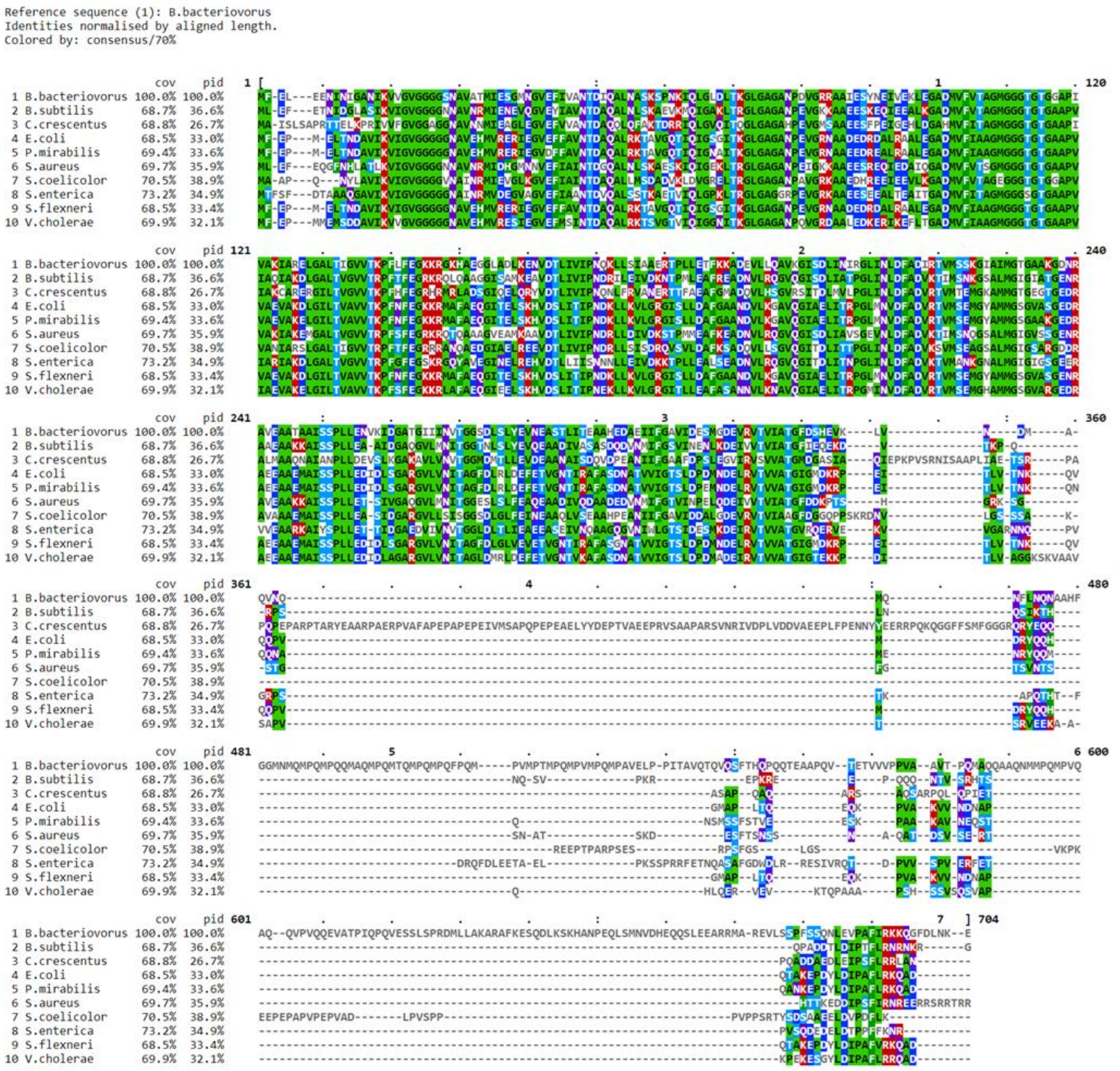


**Fig. S6. Sequence alignment of selected FtsZ proteins.** Protein sequences of 10 strains were aligned using T-Coffee algorithm (Clustal W, version 1.83) with default parameters and visualized using MView (version 1.63). FtsZ protein sequences were taken from the Uniprot database (date of access: 9 November 2022).

**A**

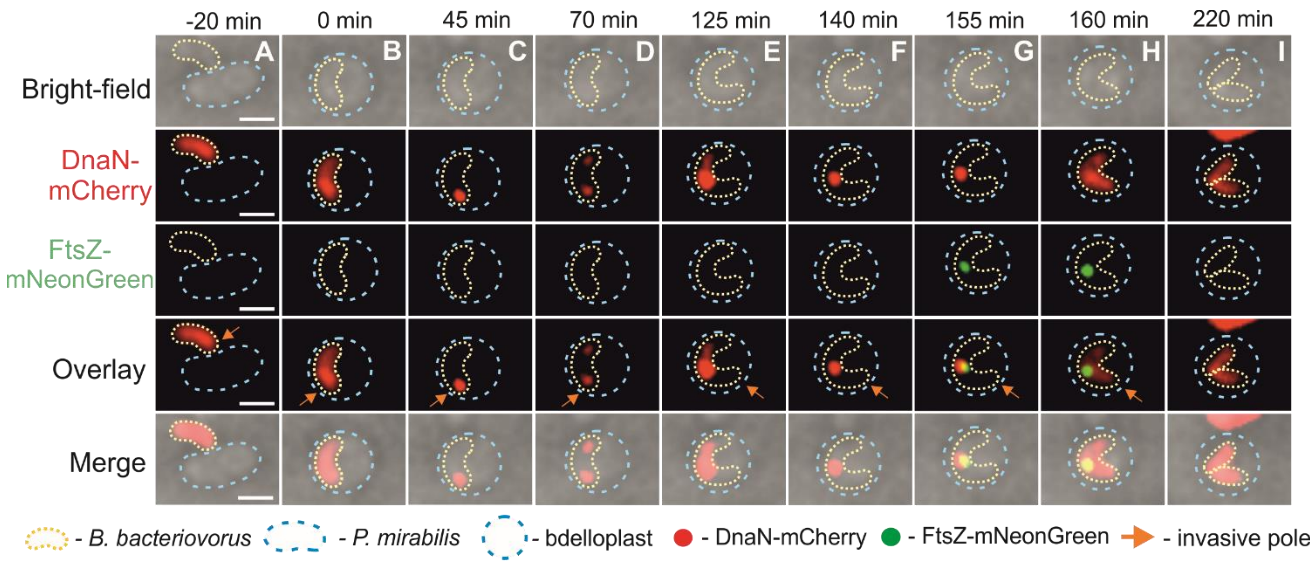


**B**

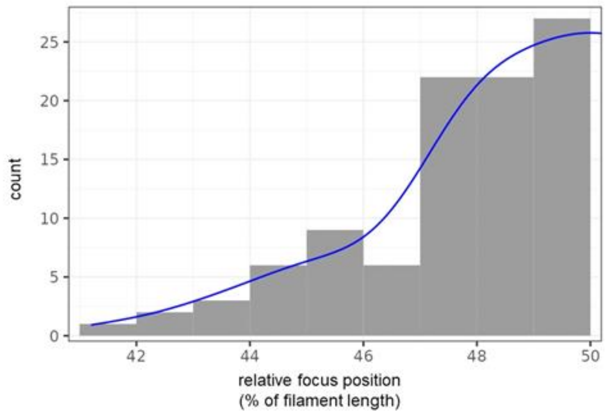


**Fig. S7. Spatiotemporal analysis of chromosome replication and filament septation in a *B. bacteriovorus* cell growing in *P. mirabilis*. (A)** The localization of replisomes (red) and divisome (green) in a predatory cell *B. bacteriovorus* growing inside the *P. mirabilis* bdelloplast. (**A**) Attachment of *B. bacteriovorus* to *P. mirabilis* cell. (**B**) Bdelloplast formation, time = 0 min. (**C**) Assembly of the first replisome at the invasive pole of *B. bacteriovorus* cell - the start of chromosome replication. (**D-F**) Further steps of chromosome replication; splitting of replication forks (the appearance of the second DnaN-mCherry focus at the 'flagellated' pole) and merging of replication forks at the midcell. (**G**) The appearance of FtsZ-mNeonGreen focus, which colocalized with the merged replisomes. (**H-I**) Termination of predatory chromosome replication (disassembly of replisomes) and filament division into two progeny cells. **(B)** Localization of divisome in relation to filament length in *B. bacteriovorus*. Subcellular localization of FtsZ-mNeonGreen foci was analyzed in cells (n=100) when the strongest fluorescent signal was observed.

Photos represent merged bright-field and fluorescence (red and green) images. The *B. bacteriovorus* cell and the bdelloplast are marked by yellow and blue dotted lines, respectively. Scale bar = 1 µm. The full time-lapses are shown in Video S7.

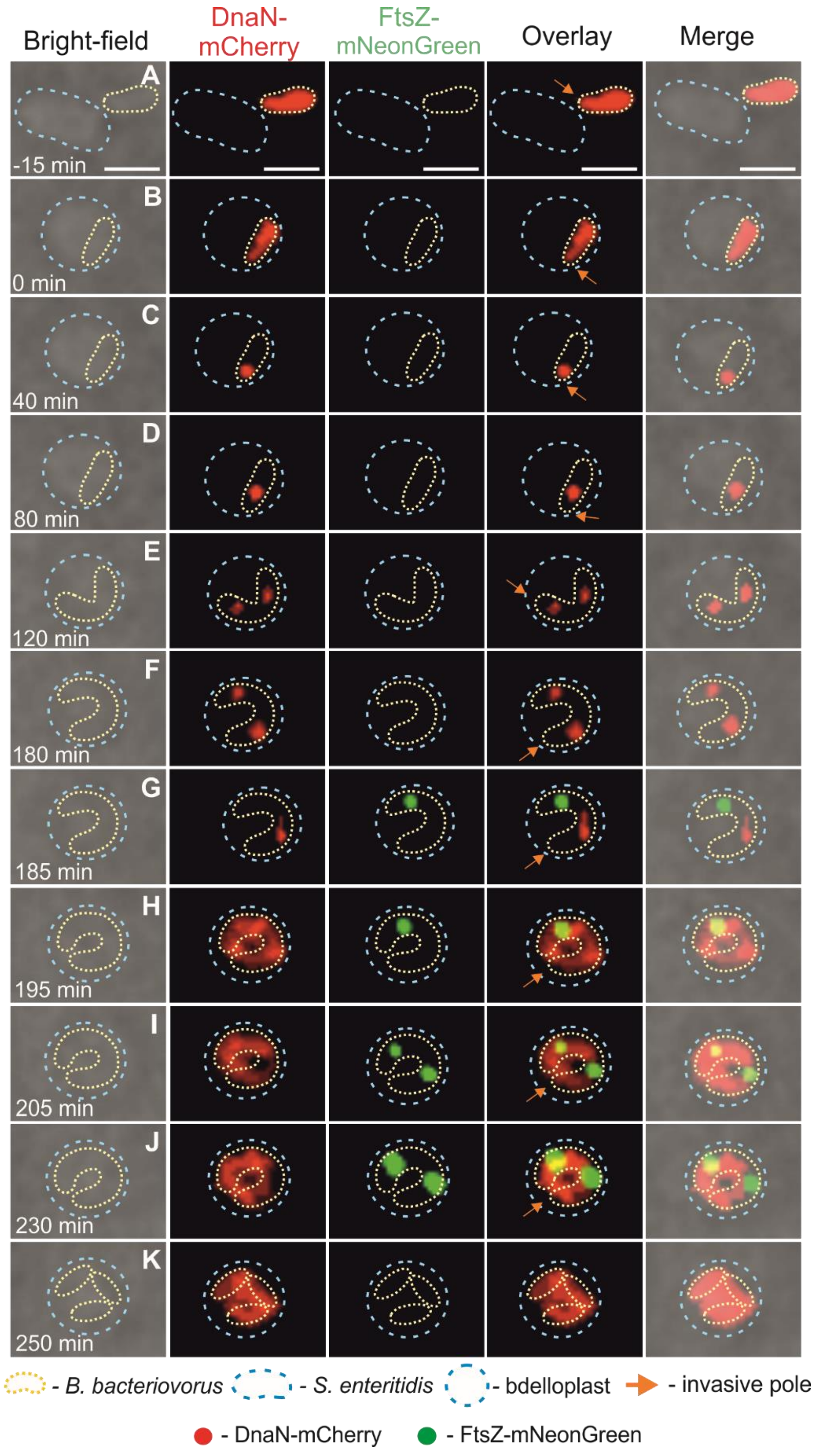


**Fig. S8. Replisome dynamics in relation to divisomes position during the cell cycle in *B. bacteriovorus* cell growing inside *S. enteritidis* and creating three progeny cells.** Time-lapse analysis of DnaN-mCherry and FtsZ-mNeonGreen signals localization during the cell cycle in *B. bacteriovorus* undergoes in *S. enteritidis*. (**A**) Attachment of *B. bacteriovorus* to a *S. enteritidis* cell. (**B**) Bdelloplast formation, time = 0 min. (**C**) Appearance of the first DnaN-mCherry focus at the invasive pole of *B. bacteriovorus* cell - initiation of chromosome replication. (**D**) Migration of DnaN-mCherry signal towards the pilus pole. (**E-F**) Reinitiation of DNA replication from the mother chromosome located at the invasive pole and migration of both replisomes toward the opposite pole. (**G**) The disappearance of one replisome located near 'flagellated' pole - termination of multiplication of one chromosome. At the same time and localization appearance of the first FtsZ-mNeonGreen foci. (**H-J**) Termination of replication process (disassembly of replisomes) and appearance of the second divisome. (**K**) Termination of septation process with formed three daughter cells.

Photos represent merged bright-field and fluorescence (red and green) images. The *B. bacteriovorus* cell and the bdelloplast are marked by yellow and blue dotted lines, respectively. Scale bar = 1 μm. The full time-lapse is shown in Video S8.

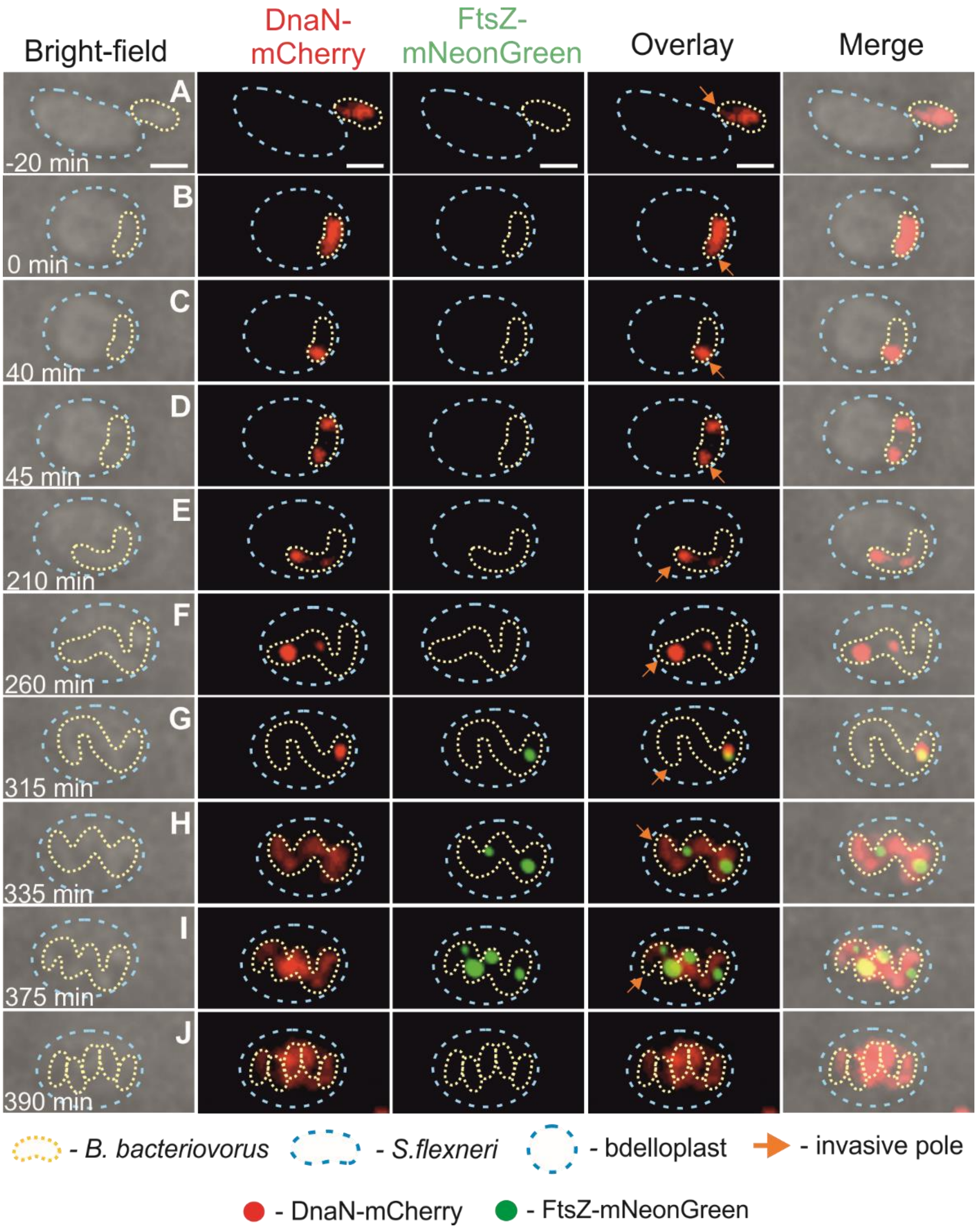


**Fig. S9. Spatiotemporal analysis of representative *B. bacteriovorus* cell showing the localization of replisomes (red) and divisomes (green) in a predatory cell growing inside the *S. flexneri* bdelloplast.** (**A**) Attachment of *B. bacteriovorus* to *S. flexneri* cell. (**B**) Bdelloplast formation, time = 0 min. (**C**) Assembly of the first replisome at the invasive pole – the beginning of multiplication DNA. (**D-F**) Migration of the first replisome toward the opposite pole and appearance of another DnaN-mCherry signals from the invasive (reinitiation of DNA replication). (**G**) Reinitiation of DNA replication from the chromosome located at the 'flagellated' pole and appearance of the first FtsZ-mNeonGreen foci localized at the same position as the last replisome. (**H-I**) Termination of DNA replication (disassembly of replisome) and appearance of another divisomes. (**J**) Termination of septation process and division into five progeny cells.

Photos represent merged bright-field and fluorescence (red and green) images. The *B. bacteriovorus* cell and the bdelloplast are marked by yellow and blue dotted lines, respectively. Scale bar = 1 µm. The full time-lapse is shown in Video S9.

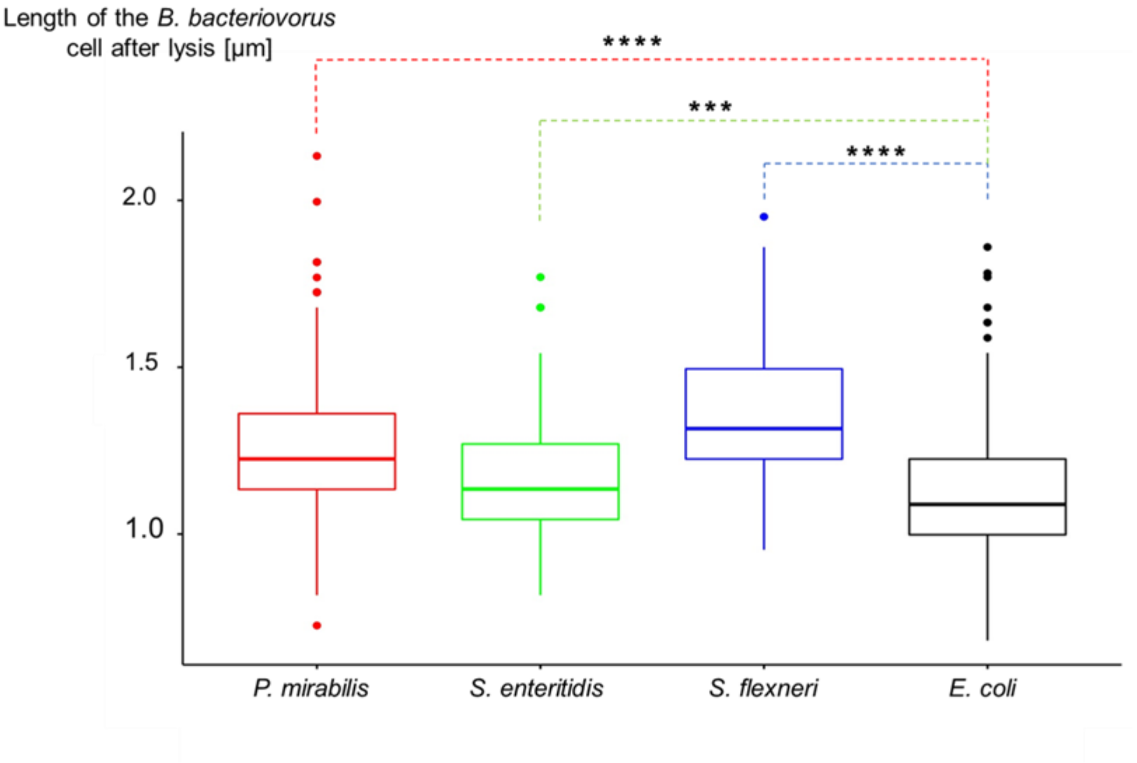


**Fig. S10. Length of mature *B. bacteriovorus* cells released from different hosts.** The average length was obtained after statistical analysis performed on 250 predator cells after bdelloplast lysis for each host and compared to the size of progeny cells released from the model host, *E. coli*. The average length of *B. bacteriovorus* cells was 1.25±0.03 µm for *P. mirabilis* (p-value<$10^{-16}$), 1.16±0.03 µm for *S. enteritidis* (p-value <$10^{-4}$), and 1.35±0.03 µm for *S. flexneri* (p-value<$10^{-16}$), while for *E. coli* was 1.11±0.04 µm. Statistical significance was assessed using the T-student test. Results distribution is shown as a boxplot with a central line depicting median value and box boundaries corresponding to the first and third quartiles. For all measured values 95% confidence interval was calculated, assuming a normal distribution of results.

**Videos:**

Video S1

Time-lapse imaging of replisomes in *B. bacteriovorus* during proliferation inside *P. mirabilis*. Subcellular localization of DnaN-mNeonGreen (green) in strain HD100 DnaN-mNeonGreen. Bright-field (grey) signals were taken every 5 min.

Video S2

Time-lapse imaging of chromosome replication and *oriC* regions segregation in *B. bacteriovorus* during proliferation inside *P. mirabilis*. Subcellular localization of DnaN-mCherry (red) and mNeonGreen-ParB (green) in strain HD100 mNeonGreen-ParB/DnaN-mCherry. Bright-field (grey) signals were taken every 5 min.

Video S3

Time-lapse imaging reflected conversion of the 'flagellated' pole into an invasive pole inherited by progeny cell of *B. bacteriovorus* during growing inside *P. mirabilis*. Localization of mNeonGreen-ParB (green) in strain HD100 mNeonGreen-ParB/DnaN-mCherry indicate the newly formed invasive pole.

Video S4

Time-lapse imaging of chromosome replication and *oriC* regions segregation in *B. bacteriovorus* cell during predation on *S. enteritidis* and formation of three daughter cells. Subcellular localization of DnaN-mCherry (red) and mNeonGreen-ParB (green) in strain HD100 mNeonGreen-ParB/DnaN-mCherry. Bright-field (grey) signals were taken every 5 min.

Video S5

Time-lapse imaging of chromosome replication and *oriC* regions segregation in *B. bacteriovorus* cell during predation on *S. enteritidis* and formation of four progeny cells. Subcellular localization of DnaN-mCherry (red) and mNeonGreen-ParB (green) in strain HD100 mNeonGreen-ParB/DnaN-mCherry. Bright-field (grey) signals were taken every 5 min.

Video S6

Time-lapse imaging of chromosome replication and *oriC* regions segregation in *B. bacteriovorus* cell during proliferation in *S. flexneri*. Subcellular localization of DnaN-mCherry (red) and mNeonGreen-ParB (green) in strain HD100 mNeonGreen-ParB/DnaN-mCherry. Bright-field (grey) signals were taken every 5 min.

Video S7

Time-lapse imaging of chromosome replication in relation to filament septation in *B. bacteriovorus* cell during growing in *P. mirabilis*. Subcellular localization of DnaN-mCherry (red) and FtsZ-mNeonGreen (green) in strain HD100 FtsZ-mNeonGreen/DnaN-mCherry. Bright-field (grey) signals were taken every 5 min.

Video S8

Time-lapse imaging of chromosome replication in relation to filament septation in *B. bacteriovorus* cell inside *S. enteritidis*. Subcellular localization of DnaN-mCherry (red) and FtsZ-mNeonGreen (green) in strain HD100 FtsZ-mNeonGreen/DnaN-mCherry. Bright-field (grey) signals were taken every 5 min.

Video S9

Time-lapse imaging of chromosome replication in relation to filament septation in *B. bacteriovorus* cell during proliferating in *S. flexneri*. Subcellular localization of DnaN-mCherry (red) and FtsZ-mNeonGreen (green) in strain HD100 FtsZ-mNeonGreen/DnaN-mCherry. Bright-field (grey) signals were taken every 5 min.
